# Synthetic Maturation of Multilineage Human Liver Organoids via Genetically Guided Engineering

**DOI:** 10.1101/2020.05.10.087445

**Authors:** Jeremy J. Velazquez, Ryan LeGraw, Farzaneh Moghadam, Yuqi Tan, Jacquelyn Kilbourne, Joshua Hislop, S Liu, Davy Cats, Susana M. Chuva de Sousa Lopes, Christopher Plaisier, Patrick Cahan, Samira Kiani, Mo R. Ebrahimkhani

## Abstract

Pluripotent stem cell (PSC)-derived organoids are emerging as novel human-based microphysiological models but display immature phenotypes with limited subsets of endothelial or stromal cells. Here we demonstrate that *in vitro* manipulation of gene regulatory networks (GRNs) in PSC-derived liver organoids selected either through computational analysis or targeted tissue design can advance tissue maturation *in vitro*. Through an unbiased comparison with the genetic signature of mature livers, we identify downregulated GRNs in fetal liver organoids compared to adult livers. We demonstrate that overexpression of *PROX1* and *ATF5*, together with targeted CRISPR-based transcriptional activation of endogenous *CYP3A4*, drives maturation *in vitro*. Single cell analyses reveal hepatobiliary-, endothelial-, and stellate-like cell populations. The engineered organoids demonstrate enhanced vasculogenesis, capture native liver characteristics (e.g. FXR signaling, CYP3A4 activity), and exhibit therapeutic potential in mice. Collectively, our approach provides a genetically guided framework for engineering developmentally advanced multilineage tissues from hiPSCs.

**HIGHLIGHTS:**

- *In vitro* tissue maturation via genetically encoded molecular programs
- Computational analysis to identify maturation transcription factors in liver organoids
- Promoting vascularization of organoids via genetically encoded molecular programs
- Single cell analysis of parenchymal and non-parenchymal cells
- Modeling of native liver functions and *in vivo* therapeutic potential

**Graphical Abstract:** 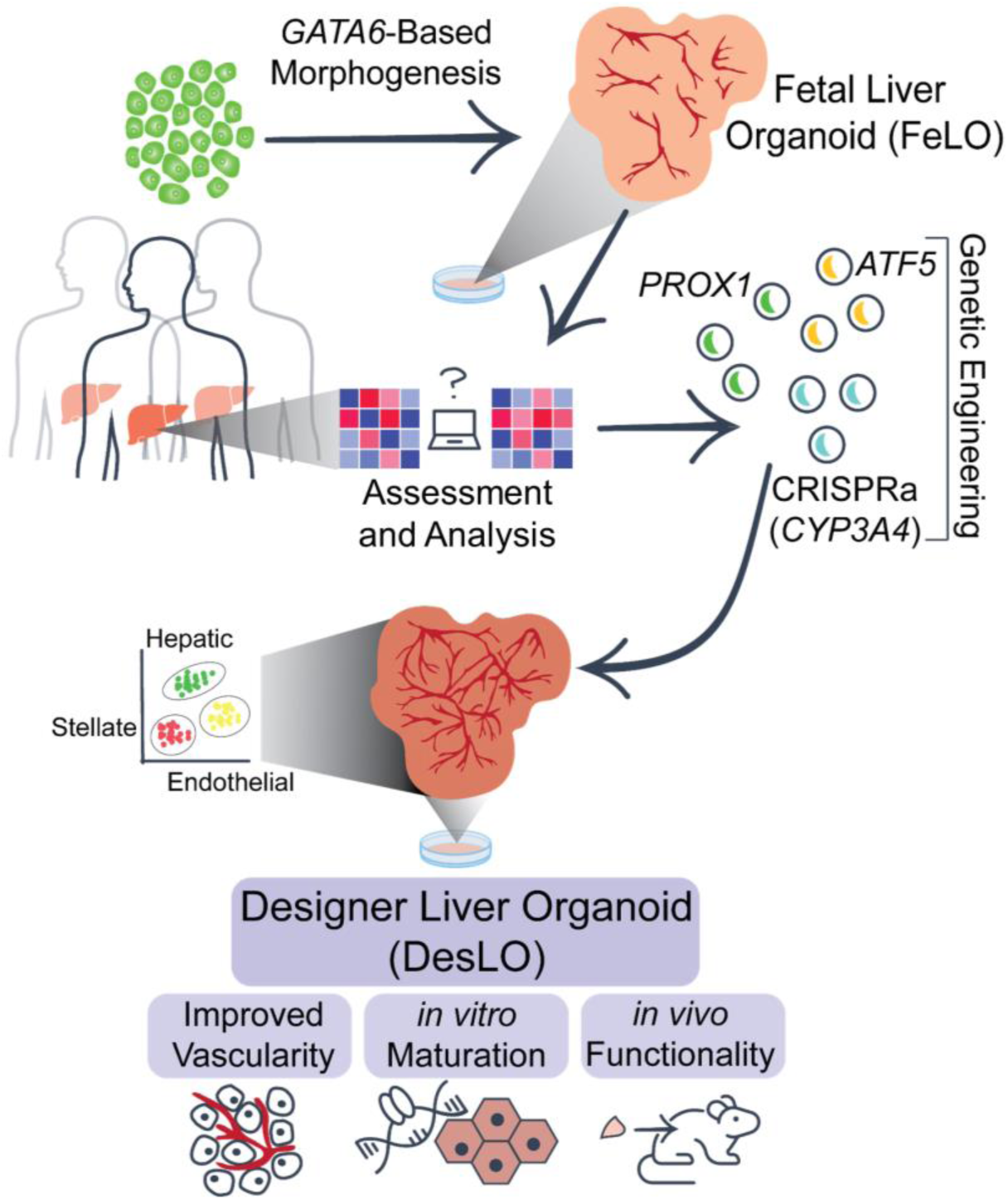

## INTRODUCTION

Self-organizing multicellular systems such as organoids have rapidly emerged at the forefront of research in biology and regenerative medicine. However, pluripotent stem cell (PSC)-derived organoids often stall developmentally and retain immature phenotypes. Organoid implantation in animal hosts can promote maturation (Takebe et al., 2013), but this process is expensive, lengthy, and lacks scalability and appropriate control over the tissue development processes. Therefore, there is a lack of rational engineering strategies to examine and direct multicellular tissues *in vitro* towards an identity close to desired adult counterparts. Most organoids developed thus far have been generated through sequential administration of growth factor cocktails in culture medium. However, this approach allows limited spatial control over target cell populations and finding an optimal combination of soluble factors that can direct and maintain multiple cell types from different germ layers is challenging. Moreover, technologies that can accelerate tissue development and bypass several months inherent to the natural process of human development will prove to be both cost effective and attractive for rapid application of organoids towards therapeutics.

Naturally evolved gene regulatory networks (GRNs) hardwired within individual cells direct cellular fates and promote collective cellular behaviors that define tissue composition, identity, and function (Bolouri and Davidson, 2002; Peter and Davidson, 2011). Here we propose a “genetically guided engineering” framework based on manipulation of cellular GRNs to direct maturation in multilineage tissues. We built upon our previously reported self-organizing “fetal” liver organoids (FeLO) which emerge from co-differentiation of progenitor cells following transient overexpression of a GATA binding protein 6 (*GATA6*) transgene in human induced PSCs (hiPSCs). This provides a human tissue that closely mimics key features of the fetal liver such as development of hepatoblasts and subsets of stromal cells. To further drive *in vitro* maturation of FeLO, we devised an approach that combines computational analysis, GRN assessment, and genetic engineering. Informed by quantitative assessment of FeLO and comparison with adult liver GRNs, we identify, build, and deliver genetic circuits that successfully trigger global tissue maturation to establish mature liver GRNs. We broadly name this rationally designed, genetically engineered FeLO as a **Designer Liver Organoid (DesLO)**. Compared to FeLO, DesLO exhibits phenotypic enhancement in several areas including development of vascular networks, bile acid sensing, and metabolic capacity. Upon implantation *in vivo*, DesLOs secrete human liver proteins and can alleviate liver disease burden. Here, we show a powerful application for engineering genetic circuits to drive maturation in human-based multicellular systems and organoids *in vitro*.

## RESULTS

### Generating a Multicellular Fetal Liver-like Organoid via *GATA6* Transposition

Previously, we employed transient and heterogeneous lentiviral expression of *GATA6* to develop FeLO from hiPSCs *in vitro* (Guye et al., 2016). Here, to refine transgene expression with efficient and stable integration and minimal toxicity, we tested a PiggyBac transposition approach. Following transient *GATA6-2A-EGFP* induction by addition of doxycycline in pluripotency medium for 5 days, we detected the emergence of endodermal (FOXA2^+^ and SOX17^+^) and mesodermal (Brachyury, T^+^) progenitors (**Figures 1A and S1A-S1D**) as expected. In the course of 15 days cells co-differentiate and self-organize into a 3-dimensional (3D) tissue with characteristics of the human fetal liver consisting of multiple cell layers spanning the surface of the tissue culture well (Guye et al., 2016). Real-time quantitative polymerase chain reaction (qPCR) confirms these data and shows downregulation of pluripotency markers *NANOG* and *SOX2* (**Figures 1B, 1C, and S1A-S1D**). Markers associated with hepatic (*HNF4A*), endothelial (*CD34*), and stromal (desmin, *DES*) cell types increase by day 17 (**Figure 1D**), suggesting steady differentiation towards a complex multicellular liver identity. Immunofluorescence staining shows polygonal alpha-1 antitrypsin^+^ (AAT) hepatocyte-like cells with E-cadherin (ECAD) expression at cell-cell junctions (**Figure 1E**). CD31^+^ and ERG^+^ endothelial cells form vascular structures associated with DES^+^ and NES^+^ stromal cells that integrate with the hepatic layer by day 15 (**Figures 1F and S1E**). Transcriptional analyses of FeLO on days 5, 10, and 17 confirm hepatic fate and enrichment of metabolic and hematopoietic pathways (**Figures S1F and S1G**). Taken together, we show development of a multicellular fetal liver-like tissue using PiggyBac-mediated transposition of *GATA6* in hiPSCs. Next, we aimed to devise an approach to advance FeLO maturation towards adult human liver.

**Figure 1.**
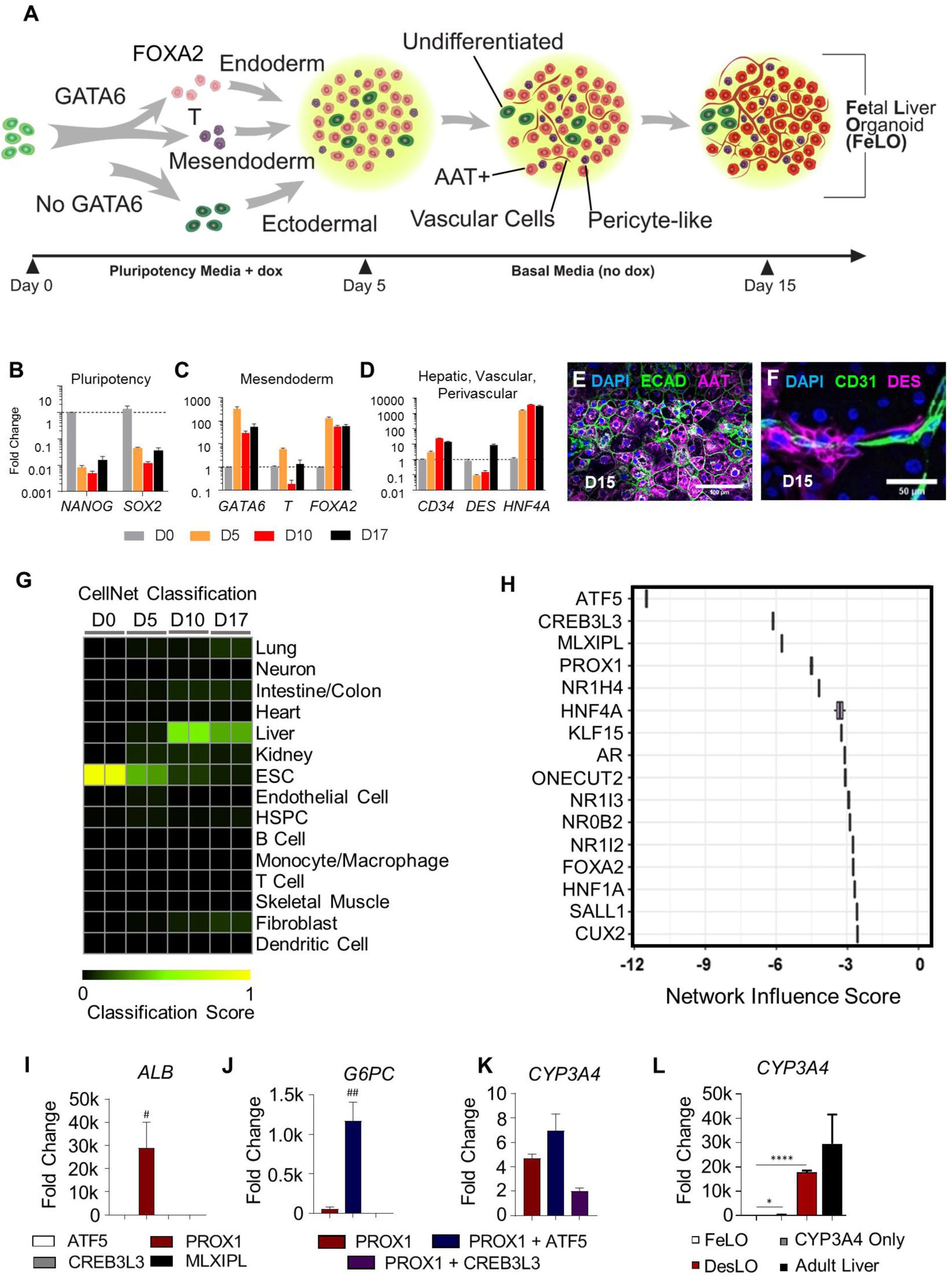
Generation and Assessment of Fetal Liver Organoid (FeLO) (A) Schematic of FeLO generation process from *GATA6*-engineered hiPSC. Full culture details are listed in the Methods section. (B-D) qPCR data of decreasing pluripotency markers (B), initial increase of early mesendoderm markers (C), and increase of hepatic (*HNF4A*), vascular (*CD34*), and stromal (*DES*), transcripts (D). Dotted lines denote reference sample expression. D0, D5, D10, and D17: day 0, 5, 10, and 17 respectively. (n=3 except n=2 for D0). (E-F) FeLO cultures stained at day 15 show ECAD tight junctions, AAT (E), DES, and CD31-expressing cells (F). (G) Heatmap showing the CellNet classification scores of FeLO for listed tissues or cell types for before induction of *GATA6* (day 0) and at day 5, 10, and 17 of culture. (H) Liver network influence score analysis using CellNet reveals transcriptional regulators that show deficiency (negative score) in day 17 FeLO compared to human liver training set. (I-L) qPCR data for transcription factor screening in FeLO, increase in *ALB* following *PROX1* induction (I), synergistic activation of *G6PC* with PROX1 and ATF5 (J), modest increase in *CYP3A4* without SynTF(*CYP3A4*) (K), and significant *CYP3A4* upregulation with SynTF(*CYP3A4*) and synergistic effect of *PROX1* and *ATF5* co-expression on *CYP3A4* activation (L). Reference sample is untransduced. # = p<0.05, ## = p<0.01 over each other condition, *p<0.05, ****p<0.0001 (n=3 except n=6 for CYP3A4 only in L). Data are represented as mean ± SEM for B-D and I-L. See also Figure S1.

### Computational Assessment of Tissue GRNs and Identification of Maturation-promoting Transcription Factors

To direct FeLO characteristics towards an adult identity, we first performed computational analysis of FeLO GRNs using CellNet (Cahan et al., 2014; Radley et al., 2017). CellNet sources RNA-Seq data from 97 studies to develop training sets, with no cell type or tissue training set using fewer than 4 distinct studies (**Figure S1H**). It compares RNA-Seq data from input samples to native cell types and tissues by both classification and GRN scores, which are measures of similarity to the target cell type or tissue and the extent to which a cell type- or tissue-specific GRN is established, respectively. 107 adult human livers from 10 studies were used to build the CellNet liver training set. Therefore, the platform offers an unbiased approach for defining tissue identity and classification of target and aberrant GRNs. CellNet analysis of RNA-Seq data from our uninduced hiPSC line (day 0) and developing tissue at days 5, 10, and 17 (FeLO) revealed high scores for embryonic stem cells (ESC) at day 0 and day 5 that steadily dropped over time concurrent with increasing liver scores (**Figures 1G and S1I**).

CellNet calculates a Network Influence Score (NIS) for transcription factors according to the degree to which their perturbation will lead to improved GRN scores. In this case, CellNet scored transcription factors according to their predicted ability to drive FeLO towards a mature human liver fate (**Figure 1H**). The top four transcription factors, whose negative score indicate under-expression relative to the liver training set (the target tissue), were activating transcription factor 5 (*ATF5*), cAMP responsive element binding protein 3 like 3 (*CREB3L3*), MLX Interacting Protein Like (*MLXIPL*), and Prospero-related homeobox1 (*PROX1*).

Focusing on these four factors, we first examined the effect of their overexpression on a panel of hepatic and endothelial markers. We devised a method to efficiently deliver expression vectors (a.k.a. gene circuits) on day 5 by dissociation to single cells followed by lentiviral transduction (see methods). Initial analysis with an mKate2 reporter showed our method yielded high transduction efficiency (**Figure S1J**). We transduced FeLO with each transgene individually, with untransduced tissue serving as a control. Analysis of gene expression by qPCR at day 17 revealed *PROX1* to substantially outperform the other factors in upregulation of a panel of mature liver genes including albumin (*ALB*), asialoglycoprotein Receptor 1 (*ASGR1*), and farnesoid X receptor (*FXR* or *NR1H4*) (**Figures 1I, S1K, and S1L**). We further tested whether expression of another factor in tandem with *PROX1* would confer any additional benefit. *PROX1* and *ATF5* co-expression augmented expression of several mature markers and displayed a strong synergistic effect on genes such as *G6PC* (**Figures 1J, 1K, and S1M**).

PROX1 and ATF5 have been shown to be involved in various facets of liver development and function. PROX1 is required for hepatocyte migration and endothelial cell development, and loss of *PROX1* leads to decreased hepatocyte counts and liver size (Kamiya et al., 2008; Seth et al., 2014). ATF5 is associated with differentiation, survival, and maturation of several cell types, including hepatocytes (Nakamori et al., 2016; Pascual et al., 2008; Wang et al., 2014). Furthermore, overexpression of the transcription factors HNF1A, HNF4A, HNF6, and CEBPA along with ATF5 and PROX1 has been used to convert fibroblasts into hepatocytes (Du et al., 2014). Given the computational assessment, initial screenings, and past literature, we chose *PROX1* and *ATF5* from our list of transcription factors.

### CRISPR-based Synthetic Transcription Factor for CYP3A4 in Liver Organoids

Overexpression of *PROX1* and *ATF5* led to a moderate induction of Cytochrome P450 3A4 (*CYP3A4*) (Figure 1K). CYP3A4 is a central hepatic enzyme responsible for metabolizing a large portion (∼30%) of clinically-used drugs (Zanger and Schwab, 2013). Achieving CYP3A4 levels comparable to adult human liver is highly sought after in PSC-derived hepatocytes for applications in drug discovery and regenerative medicine, yet previous attempts to reach this goal have been mostly ineffective (Davidson et al., 2015; Wobus and Loser, 2011). Therefore, we asked whether targeted activation of the endogenous *CYP3A4* locus could meet this need, and whether *CYP3A4* activation in parallel with PROX1/ATF5 could advance the maturity of FeLO.

We devised a synthetic transcription factor for endogenous *CYP3A4* activation (SynTF(*CYP3A4*)) based on Clustered Regularly Interspaced Short Palindromic Repeat (CRISPR)-based transcriptional activation (CRISPRa). To confine CYP3A4 activation to hepatocytes, we used a version of CRISPRa in which dCas9 is driven by an AAT promoter (pAAT) and two engineered hairpin aptamers in the gRNA recruit an MS2 bacteriophage coat protein (MCP)-fused transcriptional activation complex (Konermann et al., 2015) (**Figures S1N and S1O**). We verified the absence of known SNPs in our hiPSC cell line that could influence CYP3A4 activity (**Figure S1P**). We then designed two gRNA targeting the *CYP3A4* promoter and transduced FeLO at day 5 with all components of the CRISPRa complex. At day 17, we found significant upregulation of *CYP3A4* expression (approximately 400-fold) (**Figure 1L**). Strikingly, when we co-delivered SynTF(*CYP3A4*) with the *PROX1* and *ATF5* circuits, we discovered a dramatic synergistic effect which resulted in an almost 20,000-fold upregulation of *CYP3A4*, reaching expression levels on par with adult liver tissue (**Figure 1L**). Combined delivery of *PROX1, ATF5*, and SynTF(*CYP3A4*) to FeLO leads to a new engineered tissue that we have dubbed as DesLO.

### Genetic Manipulation with Computational Analysis Reveals Tissue Maturation

To investigate the effect of each genetic factor, we generated deconstructed variations of DesLO in parallel using our delivery method (**Figure 2A**) by introducing circuits encoding *PROX1* only, *PROX1* and *ATF5*, and the full combination of *PROX1, ATF*5, and SynTF(*CYP3A4*). Through comparison to each other and age-matched FeLO we could examine the influence of each layer of engineering on *in vitro* maturation towards the target adult liver identity. Strikingly, CellNet analyses revealed that *PROX1, ATF5*, and SynTF(*CYP3A4*) each contribute to a stepwise increase in global liver GRN and tissue classification scores (**Figures 2B and 2C**). Comprehensive gene set enrichment and pathway analyses using Enrichr also revealed increasing alignment to liver identity and associated pathways with the introduction of each circuit (**Figures S2A and S2B**). This occurred in parallel with repression of aberrant lineage expression such as intestinal marker *CDX2* and pancreas and intestinal markers somatostatin (*SST*) and pancreatic and duodenal homeobox 1 (*PDX1*) (**Figure S2C**). To cross-validate CellNet findings with an alternative methodology we used KeyGenes (Roost et al., 2015). Using human tissue training sets, KeyGenes classified DesLO as liver, demonstrating clear improvement in liver identity and noticeable elimination of the aberrant intestinal signature upon our genetic manipulation (**Figure 2D**).

**Figure 2.**
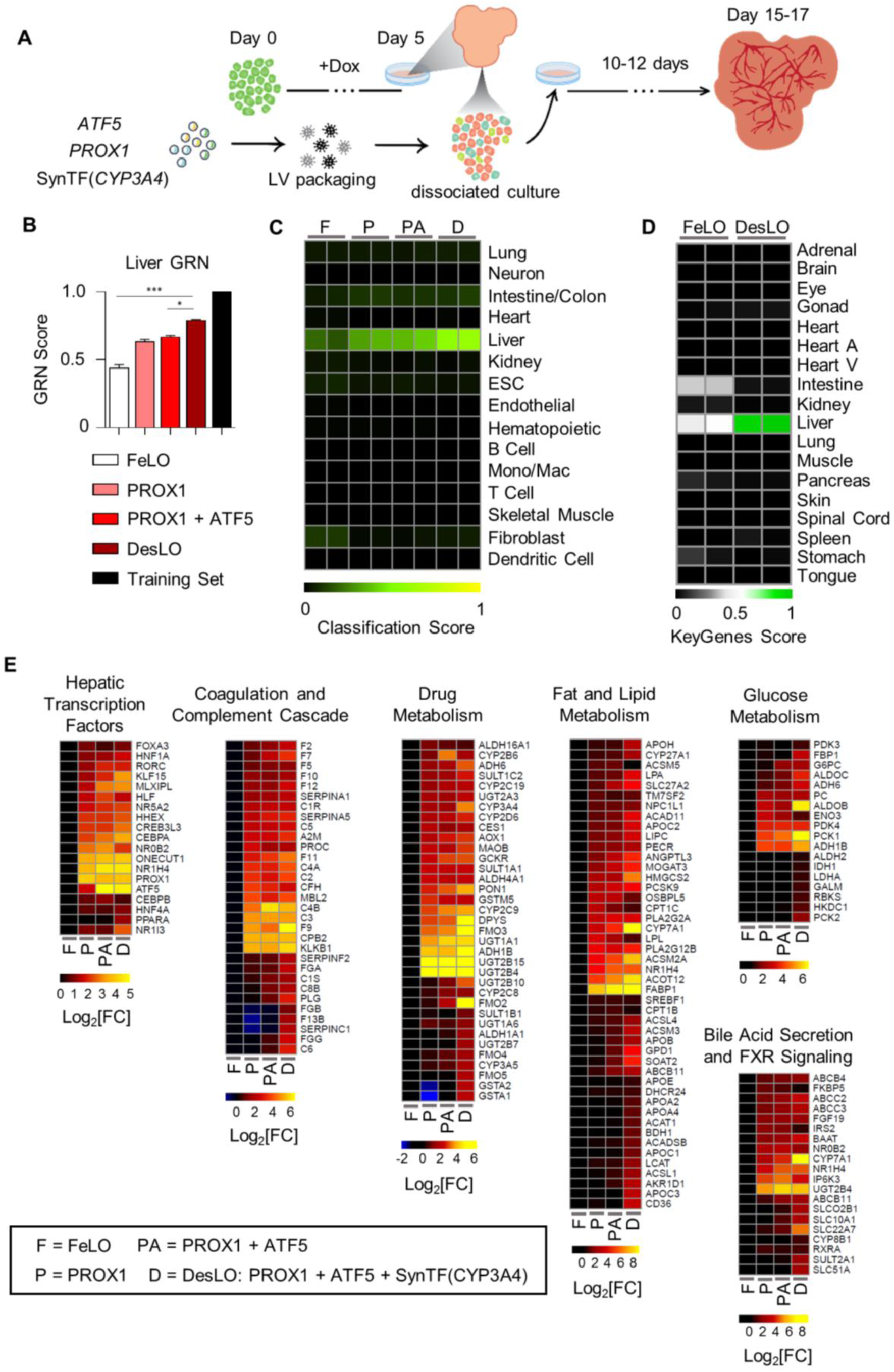
Genetic Engineering Improves Hepatic Identity and Tissue Maturation. (A) Schematic of engineering strategy using dox-inducible *GATA6* and lentiviral delivery of *ATF5, PROX1*, and SynTF(*CYP3A4*) gene circuits to develop designer liver organoids (DesLO). Full details in methods. (B) Stepwise increase in CellNet liver gene regulatory network (GRN) score by increasing the components transduced at day 5 from FeLO to DesLO in day 17 tissues. *p<0.05, ***p<0.001 (n=2). (C) Heatmap showing the CellNet classification scores for the listed tissues/organs or cell types for day 17 FeLO (F), PROX1 only (P), PROX1+ATF5 only (PA) and DesLO (D) samples (Mono/Mac: Monocyte/Macrophage). (D) Heatmap showing the KeyGenes scores for the listed tissues or cell types for day 17 FeLO and DesLO. (Heart A: Heart atrium; Heart V: Heart ventricle). (E) Heatmaps showing enrichment in pathways relative to FeLO for hepatic transcription factors, complement and coagulation cascade, drug metabolism, cholesterol and lipid metabolism, glucose metabolism, and bile acid secretion and FXR signaling by increasing the components transduced at day 5 from FeLO to DesLO in day 17 tissues. (n=2, mean shown). Data are represented as mean ± SEM for B, no SEM listed for library training set. See also Figure S2

Further examination of specific pathways critical for adult liver function showed that each layer of engineering activated a subset of liver GRNs, successively moving the synthetic tissue closer to a native liver signature. Through our intervention we observed a progressive enrichment in important pathways for glucose, lipid, cholesterol, and drug metabolism, complement and coagulation cascades, FXR signaling and bile secretion, and characteristic hepatic transcription factors (**Figure 2E**). Closer examination shows that *PROX1* induces a large portion of the increases in gene expression across these pathways, including liver transcription factors *NR1H4* (FXR), *HNF1A, ONECUT1* (HNF6), and *NR5A2* (LRH-1). Addition of the *ATF5* circuit further enriched genes across pathways such as transcription factors *MLXIPL* and *CEBPA*, the bile acid transporter *SLC10A1* (NTCP), regulator of glucose homeostasis *G6PC*, hepatokine regulator of plasma lipids angiopoietin like 3 (*ANGPTL3)*, and the enzyme *CYP2C8*, which metabolizes retinoic acid and polyunsaturated fatty acids.

Remarkably, introduction of SynTF(*CYP3A4*) in tandem with *PROX1* and *ATF5* revealed pervasive upregulation across pathways and raised global liver GRN score by ∼20% over *PROX1*/*ATF5* alone (**Figures 2B and 2E**). The presence of CYP3A4 enriched expression of transcription factors *CREB3L3, HNF4A, PPARA*, and *NR1I3* (CAR); coagulation and complement cascade components (e.g. *FGB, F7, F9*, and *C6)*; detoxification enzymes glutathione S-transferases *GSTA1* and *GSTA2*; apolipoprotein subunits; rate limiting enzyme in gluconeogenesis *PCK1*; ATP-binding cassette transporters; *ANGPTL3;* and *CYP7A1*, which metabolizes cholesterol to synthesize bile acids. Our findings warrant future investigation and may indicate novel mechanisms of liver metabolic maturation. In fact, it was shown CYP3A4 can metabolize bile acids which can then activate transcription of nuclear receptors (Bodin et al., 2005). Additionally, bile acid and derivatives promote hepatic differentiation (Sawitza et al., 2015). Thus, it is possible that CYP3A4-produced metabolites induce hepatocyte maturation via autocrine feedback loops.

### Benchmarking DesLO Against Previously Developed Cells and Tissues

To better characterize DesLO with respect to other reported PSC-derived human liver organoids (Akbari et al., 2019; Asai et al., 2017; Ouchi et al., 2019), we performed a meta-analysis of available RNA-Seq data using CellNet. We also included data for fibroblast-derived induced hepatocytes from Du et al., 2014, along with primary hepatocytes and liver tissue from two independent studies as positive controls (**Figures 3A and 3B**). These controls showed high GRN and classification scores, confirming the robust network identification by CellNet. Our analysis revealed DesLO to have the highest liver GRN and classification scores among all organoid samples, with only cryopreserved PHH, fresh PHH, and whole liver tissue scoring higher. DesLO achieved an approximately 50% higher GRN score than the next highest scoring organoid. In the classification heatmap, native human liver showed no apparent signatures from other organs. DesLO also had minimal aberrant signatures while some other tested samples showed residual signatures associated with tissues such as intestine/colon, fibroblasts, or kidney. Liver bud samples (day 2 and 6) from Asai et al. showed low liver classification scores but they are prior to implantation, and *in vivo* maturation would be expected to occur when implanted (Asai et al., 2017; Takebe et al., 2013). Collectively our analysis indicates that, at the level of tissue classification and GRN state, DesLO developed in this study reach a level of maturation not attained by past efforts assessed here.

**Figure 3.**
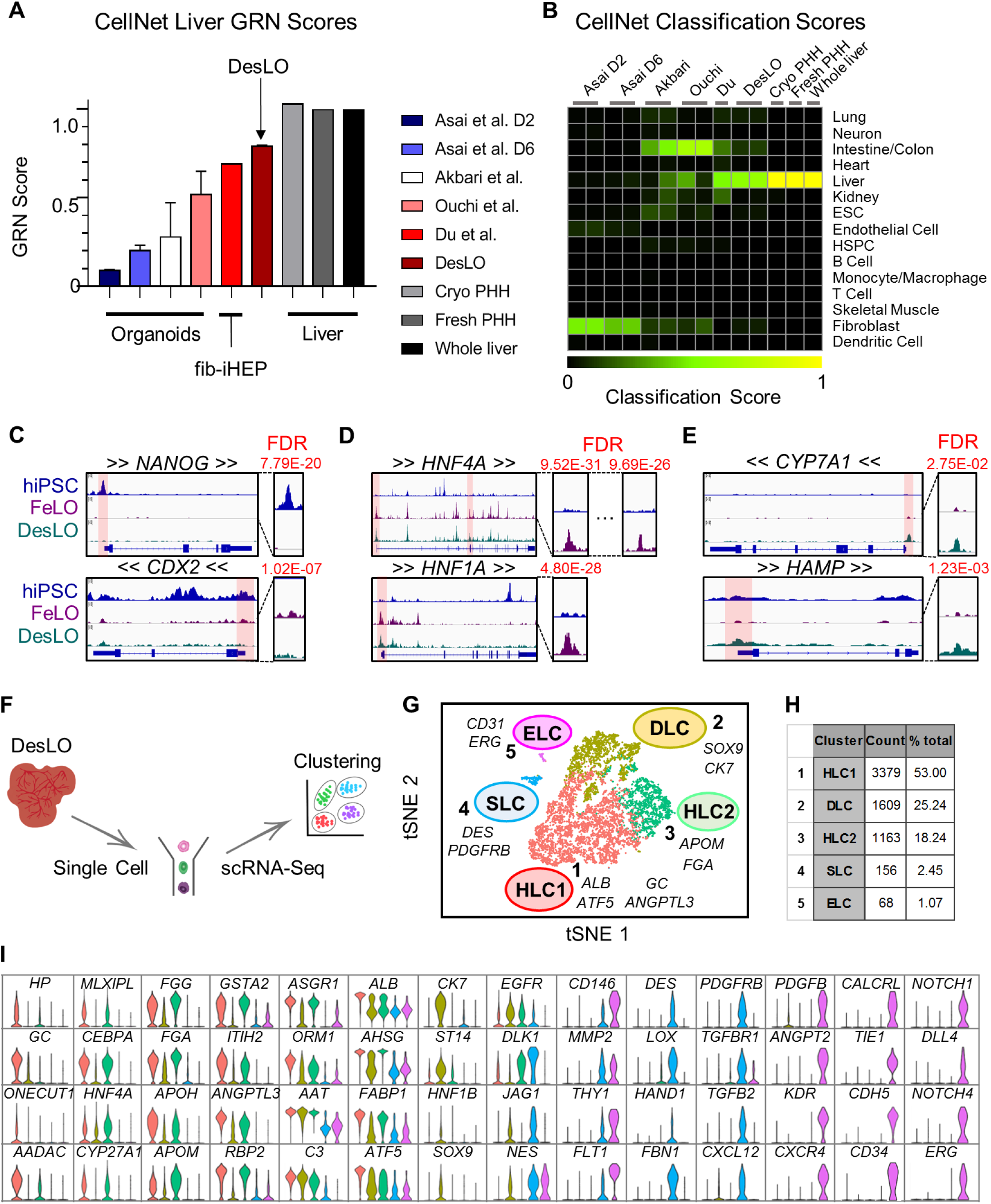
Cross-study Comparison of *in vitro*-derived Organoids, ATAC-Seq, and scRNA-Seq Analysis of DesLO. (A) CellNet meta-analysis of organoids showing GRN scores for previously reported select liver organoids from hiPSCs, fibroblast-derived induced hepatocytes (fib-iHEP), cryopreserved primary human hepatocytes (Cryo PHH), freshly isolated primary human hepatocytes (Fresh PHH), and whole liver tissue. All samples are n=2 except for Du et al., Fresh PHH, Cryo PHH, and Whole liver samples which are n=1. (B) CellNet meta-analysis of organoids showing classification scores determined for previously reported select liver organoids from hiPSCs, fib-iHEP, Cryo PHH, Fresh PHH, and whole liver tissue. (C) ATAC-Seq results showing loss of chromatin accessibility on *NANOG* and *CDX2* between hiPSCs, FeLO, and DesLO. FDR of differential peak comparison is indicated for the zoomed promoter regions highlighted in transparent red. (D) ATAC-Seq results showing increases in chromatin accessibility in *HNF4A* and *HNF1A* between hiPSCs and FeLO. FDR of differential peak comparison is indicated for the zoomed promoter regions highlighted in transparent red. (E) ATAC-Seq results showing increases in chromatin accessibility in *CYP7A1* and *HAMP* between FeLO and DesLO. FDR of differential peak comparison is indicated for the zoomed promoter regions highlighted in transparent red. (F) Schematic of single cell analysis pipeline used for day 17 DesLO and FeLO samples. (G) t-distributed Stochastic Neighbor Embedding (tSNE) plot of day 17 DesLO scRNA-Seq data analyzed using Seurat. Cluster numbers, names, and examples of enriched genes are displayed. HLC: hepatocyte-like cells, SLC: stellate-like cells, ELC: endothelial-like cells, DLC: ductal-like cells. (H) Table showing the number of cells and percentage of total population in each DesLO cluster. (I) Violin plots showing hepatocyte-specific genes upregulated in HLC, ductal-enriched genes upregulated in DLC, stellate-specific genes upregulated in SLC, and endothelial-specific genes upregulated in ELC clusters. See also Figure S3.

### ATAC-Seq Analysis Shows Chromatin Remodeling During Formation of FeLO and DesLO from hiPSCs

We employed assay for transposase-accessible chromatin sequencing (ATAC-Seq) to epigenetically survey genomic loci enriched for open chromatin, which denotes sites associated with active transcription factor binding. An overall analysis of significant peaks indicated similarity between the FeLO and DesLO samples, which both contrasted significantly with the signature of the hiPSC samples (**Figure S3A**). Importantly, differentiation from hiPSC to FeLO captures the closing of chromatin at the promoter regions of pluripotency genes such as *NANOG* and *OCT4* (official gene symbol *POU5F1*) (**Figures 3C and S3B**). Sites associated with the aberrant intestinal lineages such as the *CDX2* promoter were also significantly decreased in accessibility, further supporting the lineage specificity and stability of the system (**Figure 3C**). Conversely, promoter regions for liver identity specification such as *HNF4A* (at both of its well-known promoters), *HNF1A, FOXA2, NR5A2*, and *ATF5* were opened in FeLO and DesLO (**Figures 3D and S3B**). This coincides with enhanced promoter-localized peaks in functional liver genes for glucose metabolism (*G6PC*), complement cascade (*C3*), lipoprotein metabolism (*APOA2*), and bile acid regulation (*NR0B2*) after differentiation to FeLO and DesLO (**Figure S3B**). Furthermore, generation of DesLO significantly increases accessibility at promoter regions of loci such as *CYP7A1, HAMP*, and *INHBE*, which are important nodes for bile acid synthesis, iron homeostasis, and activin signaling in the liver respectively (**Figures 3E and S3B**). The epigenetic trends reported here corroborate our initial findings from RNA-Seq analysis and support the existence of a robust liver identity programmed at the level of chromatin states in FeLO and DesLO (**Figures S2C, 3B, and 3C**). It is important to note that the dynamic interplay of chromatin states with transcription factor availability and concentration eventually dictates final GRN structure and liver tissue identity.

### Single Cell RNA-Seq in DesLO Confirms Heterotypic Cell Types and Hepatic Maturation

Having shown maturation in bulk DesLO tissue, we next examined the composition of FeLO and DesLO at the single cell level. We performed single cell RNA sequencing (scRNA-Seq) of day 17 FeLO and DesLO cultures (**Figure 3F**). DesLO shows a visible increase in cells positive for liver transcription factors *FOXA3, MLXIPL, CREB3L3, NR0B2, NR1H4*, and *AHR*; signaling molecules *VEGFA* and *INHBE*; functional genes regulating lipid, cholesterol, and bile acid metabolism (*ANGPTL3, ANGPTL8, APOH, CYP7A1, CYP27A1, NR0B2*, and *NR1H4*); acute phase proteins (*ALB, ORM1, ORM2, HP, C3, F2*); and gluconeogenesis regulators (*G6PC* and *PCK1*) (**Figure S3C**). Notably, expression of *DLK1*, which is known to mark hepatocytes and stellate cells during development (Huang et al., 2019), decreases significantly from FeLO to DesLO indicating the transition from a fetal signature.

Our scRNA-Seq analysis pipeline (Butler et al., 2018) identified five distinct clusters (**Figures 3G and 3H)**. Clusters 1 and 3 both show enrichment in well-known hepatic genes such as *CEBPA, HNF4A MLXIPL, ASGR1*, and *AHSG* but also differ in a subset of other genes. For instance cluster 3 expresses higher levels of *GSTA2* and *FGG* and maintains some expression of *DLK1*, whereas cluster 1 was enriched in genes such as *ALB*, haptoglobin (*HP*), GC vitamin D binding protein (*GC*), and *ANGPTL3* (**Figure 3I**). These clusters, which align strongly with bulk liver tissue (**Figure S3D**), represent the pool of hepatocyte-like cells (HLC1 and HLC2) in DesLO.

Cluster 2 had some expression overlap with HLC1 (cluster 1) and HLC2 (cluster 3) but lacked comparable levels of several hepatic genes, most notably *CEBPA*, which is known to repress differentiation of hepatoblasts towards a biliary fate (Si-Tayeb et al., 2010a) (**Figures 3I and S3E**). Conversely, the cells were enriched for biliary markers *CK7* and *SOX9* (**Figures 3I and S3E**) and express higher *ST14*, a positive marker for the clonogenic subset of cholangiocytes (Li et al., 2017). This cluster could represent a pool of biliary progenitors or ductal-like cells (DLC) (**Figures 3I and S3E**).

The endothelial-like cell (ELC) cluster 5 was then identified by expression of genes including vascular endothelial growth factor receptor 1 (*VEGFR1* or *FLT1*), VE-cadherin (*CDH5*), *CD34, ERG*, and *CD31* (**Figures 3I**). The *DES*^+^/*PDGFRB*^+^ cluster 4 expressed stellate cell markers delineated in a recent single cell report on primary isolated liver samples (**Figures 3I and S3F**) (MacParland et al., 2018). These stellate-like cells (SLCs) were enriched in *DES* and structural genes including collagen type I alpha 1 chain (*COL1A1*), transgelin (*TAGLN*), and smooth muscle actin (*ACTA2*) (**Figure 3I**). Clustering of DesLO ELC- and SLC-upregulated genes with overlapping primary liver stellate and endothelial cell scRNA-Seq data from the primary liver single cell dataset also grouped DesLO ELCs closest to zone 1 liver sinusoidal endothelial cells (LSECs) and SLCs closest to stellate cells (**Figure S3F**) (MacParland et al., 2018).

We also examined transcript levels of the genes associated with the genetic circuits used in our study. As confirmation that our *GATA6-2A-EGFP* expression circuit is not active after removal of doxycycline (a.k.a. leaky) we were unable to detect GFP expression (**Figure S3G**). Interestingly however, we found that DesLO established endogenous *GATA6* expression in the hepatobiliary and stellate clusters in line with past reports (**Figure S3H**) (De Franco et al., 2013; Liu et al., 2020). In fact, a recent study has identified a key role for GATA6 in regulation of mouse and human stellate cell lineages (Liu et al., 2020). scRNA-Seq shows that *PROX1* and *ATF5* are relatively enriched in hepatobiliary populations (**Figure S3H**). Levels of these genes further vary among the hepatobiliary clusters, implying more complex interactions shaping identity that require future investigation. To further interrogate the roles of PROX1 and ATF5 in development of hepatic versus endothelial identities, we performed magnetic bead isolation of CD34^+^ ELCs and CD34^-^/CD146^-^ hepatobiliary cells (**Figure 3I**). FACS analysis of the populations confirmed that transduction efficiency is equal among cell types (**Figure S3I**). Using qPCR to assay the genomic integration of transgene DNA, we determined that the hepatic population had significantly higher integration of *PROX1* (**Figure S3J**). FACS further showed that higher transgene levels increase the proportion of hepatic cells, corroborating the notion that differential integration rates can influence hepatic cell fate selection (**Figure S3K**).

### *In Vitro* Analysis of DesLO Shows Cellular Organization and Hepatic Functions

We first assessed DesLO via immunofluorescence staining and imaging at day 17 to confirm the presence of the hepatic, stellate, and endothelial compartments. Phase imaging of the cultures revealed polygonal morphology and tight junctions between hepatic cells with some vascular structures (**Figure 4A**). These formations were corroborated by AAT and CD34 antibody staining (**Figure 4B**). Staining further showed CK18 (KRT18), CEPBPA, and HNF4A expression in the HLCs and vascular network forming CD31-, CD34-, ERG-, and CD146-expressing ELCs with DES- and NES-expressing SLCs lining the endothelium (**Figures 4C-H, S4A, and S4B**). We also found NES expression in a subpopulation of ELCs, corroborated by our scRNA-Seq analysis and in line with the suggested role for NES in endothelial cells during angiogenesis (**Figures 3I and S4B**) (Mokry and Nemecek, 1998).

**Figure 4.**
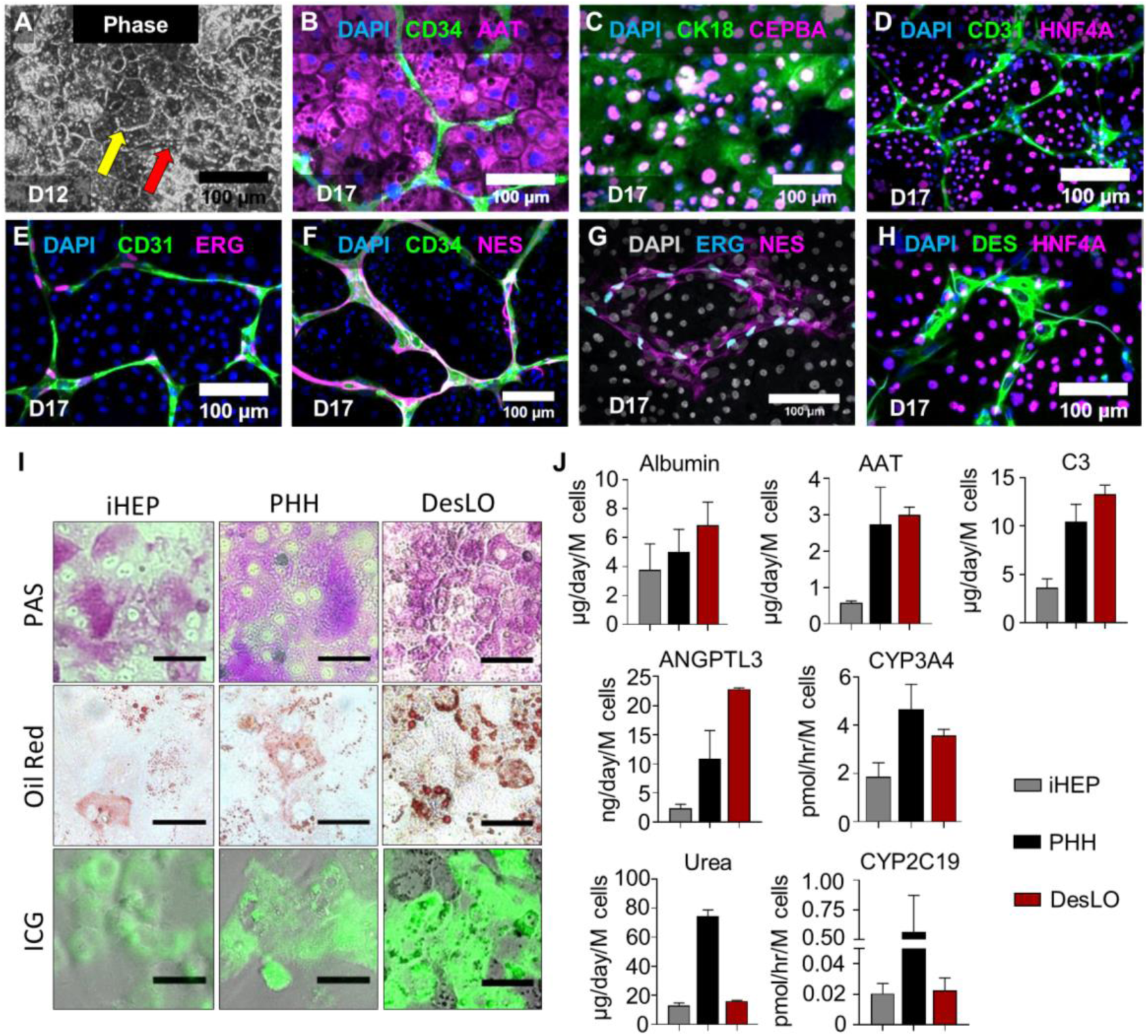
Immunofluorescence and Functional Comparison of DesLO, iHEP, and PHH cultures. (A) Phase imaging of day 12 DesLO cultures shows clear formation of tight junctions (indicated by yellow arrow) characteristic of hepatic cells in culture and visible vascular structure (indicated by red arrow) (D12: day 12). (B-H) Immunofluorescence staining for DAPI, CD34, AAT, CK18, CD31, HNF4A, ERG, NES, and DES of day 17 DesLO cultures confirming the presence of HLC, SLC, and ELC populations through staining of cell-type indicative markers (D17: day 17). (I) Periodic acid-Schiff (PAS), Oil Red O, and indocyanine green (ICG) stains indicating glycogen storage, lipid production, and solute transport activity respectively in iHEPs, PHH, and DesLO cultures. Scale bar: 50 μm. (J) ELISAs of liver-specific proteins, CYP3A4 enzymatic activity, and urea production measured in iHEP (n=6-9 from 2-3 lots), PHH (n = 6-12 from 3-4 lots except in CYP2C9 n = 3 from 1 lot), and DesLO cultures (n=3). Data are represented as mean ± SEM for J. See also Figure S4.

We next performed functional assessment of DesLO using several key hepatic metrics informed by past literature (Huch et al., 2015; Si-Tayeb et al., 2010b; Takebe et al., 2013; Zhu et al., 2014) with primary human hepatocytes (PHH) and commercially available hiPSC-derived hepatocytes (iHEPs) as controls. We performed staining for glycogen storage (Periodic acid-Schiff, PAS), lipid accumulation (Oil red O), and solute transport (Indocyanine Green, ICG) in iHEP, PHH, and DesLO, which revealed DesLO to perform similarly to PHH for each metric. (**Figure 4I**). We then quantified a panel of secreted proteins in culture media known to be important for liver function (**Figure 4J**). In DesLO we detected PHH levels of albumin and AAT. DesLO and PHH also produced comparable levels of ANGPTL3 and complement 3 (C3), the latter of which is a central component of the complement activation cascade and innate immunity produced by the liver (Sarma and Ward, 2011). Secretion of the proteins was detected in iHEPs for all measures but at lower levels than DesLO and PHH. Importantly we show enzymatic activity of CYP3A4 on par with PHH, demonstrating the functional utility of our CRISPRa approach (**Figure 4J**). Urea synthesis in both DesLO and iHEPs was detected but unable to match levels generated by PHH (**Figure 4J**). We also tested CYP2A19 activity, another drug metabolizing enzyme responsible for metabolism of ∼7% of clinically-used drugs, but observed low activity relative to PHH. Collectively we demonstrate functional capacity in several areas comparable to native PHH, but also highlight aspects to improve in future efforts.

We also tested whether DesLO can be passaged and cryopreserved, characteristics that enable usage across laboratories following generation. We show that after passaging and cryopreservation DesLO retains hepatic metrics such as production of AAT and fibrinogen on day 17 (**Figure S4C**). Additionally, tissues re-establish their vascular structures (**Figure S4D**). DesLO can also be induced to grow in different conformations based on seeding substrate, providing another dimension of control for engineering applications. A time lapse of DesLO grown on a 3D Matrigel surface demonstrates contraction into a condensed tissue (**Figure S4E**) that retains its vascular network (**Figure S4F**).

### Modeling Farnesoid X Receptor (FXR)-Mediated Responses in DesLO

For further functional analysis we sought to test the dynamic of evolved GRNs conferred by our engineering in DesLO. Activation of FXR signaling in the liver maintains bile acid homeostasis, inhibits inflammation, and controls metabolic hemostasis (Rizzo et al., 2005). It has been shown that bile acids generated in the liver can activate the FXR pathway; bile acid binding to FXR results in the secretion of fibroblast growth factor 19 (FGF19), which binds to the FGFR4-beta Klotho complex and represses CYP7A1 metabolism of cholesterol in a small heterodimer partner (SHP, official symbol NR0B2)-dependent manner (Holt et al., 2003; Inagaki et al., 2005) (**Figure 5A**).

**Figure 5.**
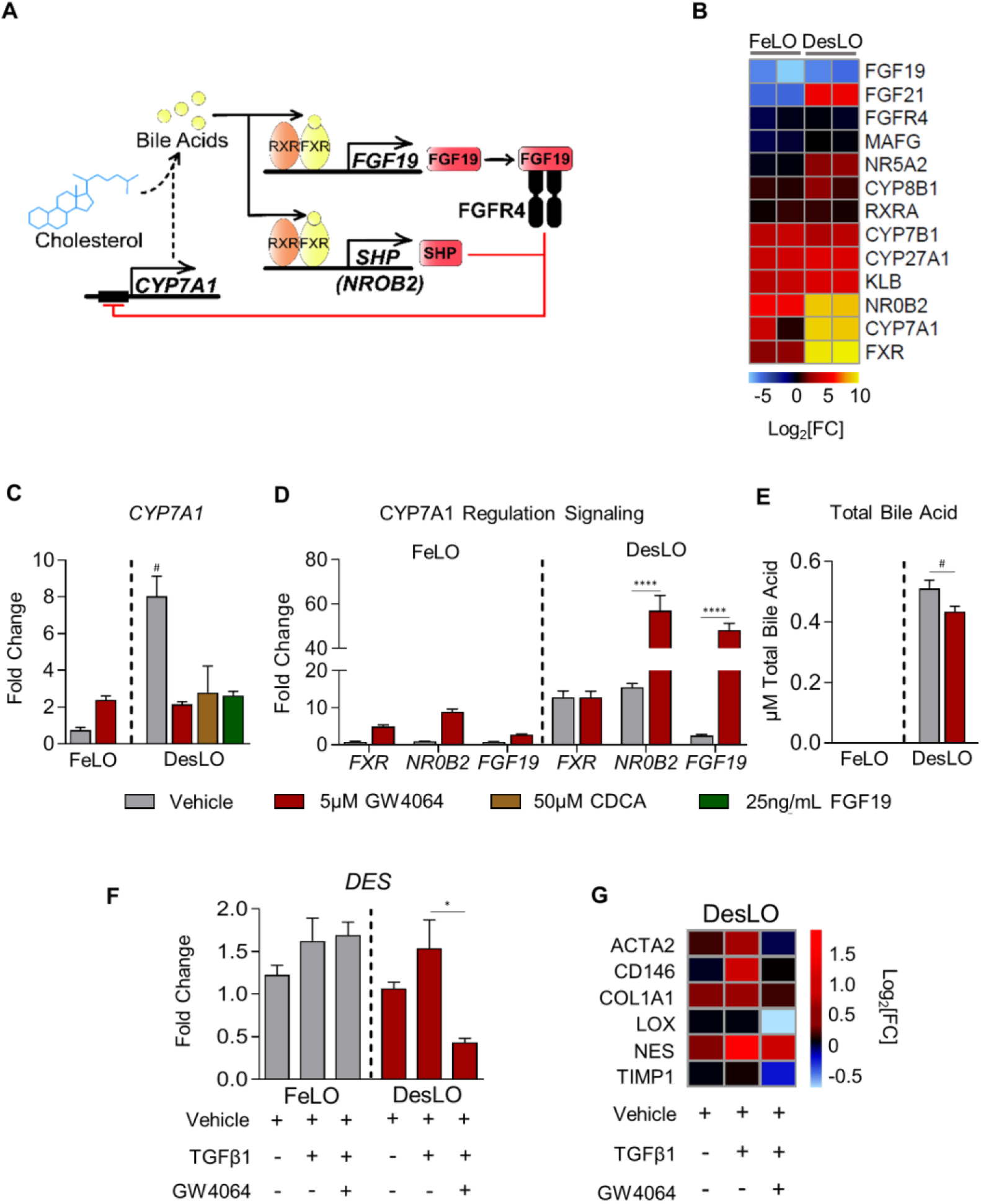
Modeling FXR Signaling and Stellate Cell-like Function in DesLO. (A) Schematic of FXR regulation of *CYP7A1* expression and bile acid synthesis. (B) Heatmap of gene expression in control FeLO and DesLO from RNA-Seq data highlighting increased DesLO expression of genes involved in FXR signaling. log2[FC] is shown in comparison to day 0 hiPSC control. (C) qPCR showing relative expression of *CYP7A1* in FeLO and DesLO with indicated additions of GW4054, CDCA, and FGF19. Fold change is over FeLO vehicle. Significance of DesLO vehicle is relative to all other DesLO, # = p<0.05 (n=3). (D) qPCR showing expression of *FXR, NR0B2, FGF19* in FeLO and DesLO vehicle controls and with addition of GW4064. Fold change is over FeLO vehicle. Significance relative to DesLO vehicle control for each target. ****p<0.0001 (n=3). (E) Total bile acid in FeLO and DesLO with addition of GW4064. Significance relative to DesLO vehicle control. #p<0.05, one-tailed t-test (n=3). (F) qPCR for *DES* (Desmin) in FeLO and DesLO with addition of TGFβ1 and GW4064. Significance relative to DesLO vehicle control. *p<0.01 (n=3). (G) Heatmap representation of qPCR data of fibrosis-associated genes from DesLO samples dosed with TGFβ1 or TGFβ1 and FXR agonist GW4064 compared against non-dosed controls. Values were normalized by median expression of DesLO vehicle control (n=3). Data are represented as mean ± SEM for C-F. See also Figure S5.

We found key mediators of the FXR/bile acid synthesis regulatory pathway upregulated in DesLO including *CYP7A1, FXR*, and *NR0B2* (**Figures 2E and 5B**). We therefore investigated whether the activation of this network provides the capacity to model the physiological dynamics of the bile acid synthesis pathway. qPCR confirmed increased expression of *CYP7A1* in DesLO. Upon addition of a potent synthetic FXR agonist, GW4064, *CYP7A1* was repressed 4-fold in DesLO, but was not repressed in FeLO cultures (**Figure 5C**). DesLO was further tested with addition of the natural bile acid chenodeoxycholic acid (CDCA) and FGF19 to the media, which also resulted in the expected downregulation of *CYP7A1* (**Figure 5C**). When compared to PHH and iHEPs, DesLO mirrored the response of PHH, while iHEPs and FeLO were unable to recapitulate this effect (**Figure S5A**).

To further interrogate the dynamics of this network we surveyed *FXR, NR0B2*, and *FGF19* expression following GW4064 treatment. DesLO displayed a potent upregulation of *NR0B2* and *FGF19* (**Figure 5D**). Such events, also observed in PHH, were less pronounced in FeLO and iHEP cultures, further supporting the presence of GRNs responsive to environmental perturbations in DesLO and PHH (**Figures 5D and S5B**). Finally, an assay of total bile acid in culture media revealed that while FeLO was unable to generate detectable amounts of bile acid from cholesterol in the medium, DesLO was capable of both producing bile acid and successfully recapitulating the role of FXR activation in repression of bile acid synthesis upon addition of GW4064 (**Figure 5E**).

FXR agonists are being tested in clinical trials for controlling liver inflammation, non-alcoholic fatty liver disease, and fibrosis (Han, 2018). Since DesLO contains a population of SLCs, we asked whether SLCs can sense and respond to fibrotic and FXR-mediated antifibrotic stimuli. We induced fibrosis via transforming growth factor beta-1 (TGFβ1) supplementation to the culture medium (Coll et al., 2018) with and without FXR activation. Staining showed DES^+^ SLCs expand relative to the vehicle control (**Figure S5C**) with qPCR corroborating the increase in *DES* levels (**Figure 5F**). Interestingly, while FeLO and DesLO both responded modestly to TGFβ1 treatment, only DesLO could reliably capture the antifibrotic effect of FXR activation (**Figure 5F**). We also assessed an array of genes relevant to the anti-fibrotic response in DesLO by qPCR. Our analysis revealed that DesLO upregulates fibrogenic genes such as *COL1A1 and ACTA2*, which are downregulated in response to the FXR agonist (**Figure 5G**). Collectively the data suggest potential for DesLO and its SLCs in modeling fibrogenic and anti-fibrogenic effects.

### Augmented Vascular Network Derived by Genetic Engineering

To further investigate the non-parenchymal cells in DesLO, we examined ELCs and development of vascular networks. Time lapse analyses of FeLO endothelial cell GRN status revealed a small but significant increase in GRN score in DesLO (**Figure S6A**). This finding is in line with the native liver GRN score and above the hepatocyte-only sample (**Figure S6A**). *PROX1* overexpression induced many key genes associated with active angiogenesis, which were maintained or enriched by introduction of *ATF5* and SynTF(*CYP3A4*) gene circuits (**Figure 6A)**. We found upregulation of key growth factors and receptors such as *VEGFA*, placental growth factor (*PGF*), and kinase insert domain receptor (*KDR*), along with endothelial identity markers such as *ERG, CD34*, and vascular endothelial cadherin (*CDH5*). Next, we performed image analysis of vascular networks. DesLO contained a vast, interconnected network of vasculature whereas FeLO was more sparse (**Figures 6B and 6C**). Analysis on days 11, 14, and 17 revealed that the total vessel length, vessel percentage area, and number of vascular junctions were increased in DesLO relative to control FeLO (**Figures 6D and S6B)**. Additionally, vessel metrics showed decreasing trends in FeLO from day 14 to 17, indicating instability of the vascular network, while the same measurements for DesLO cultures remained stable.

**Figure 6.**
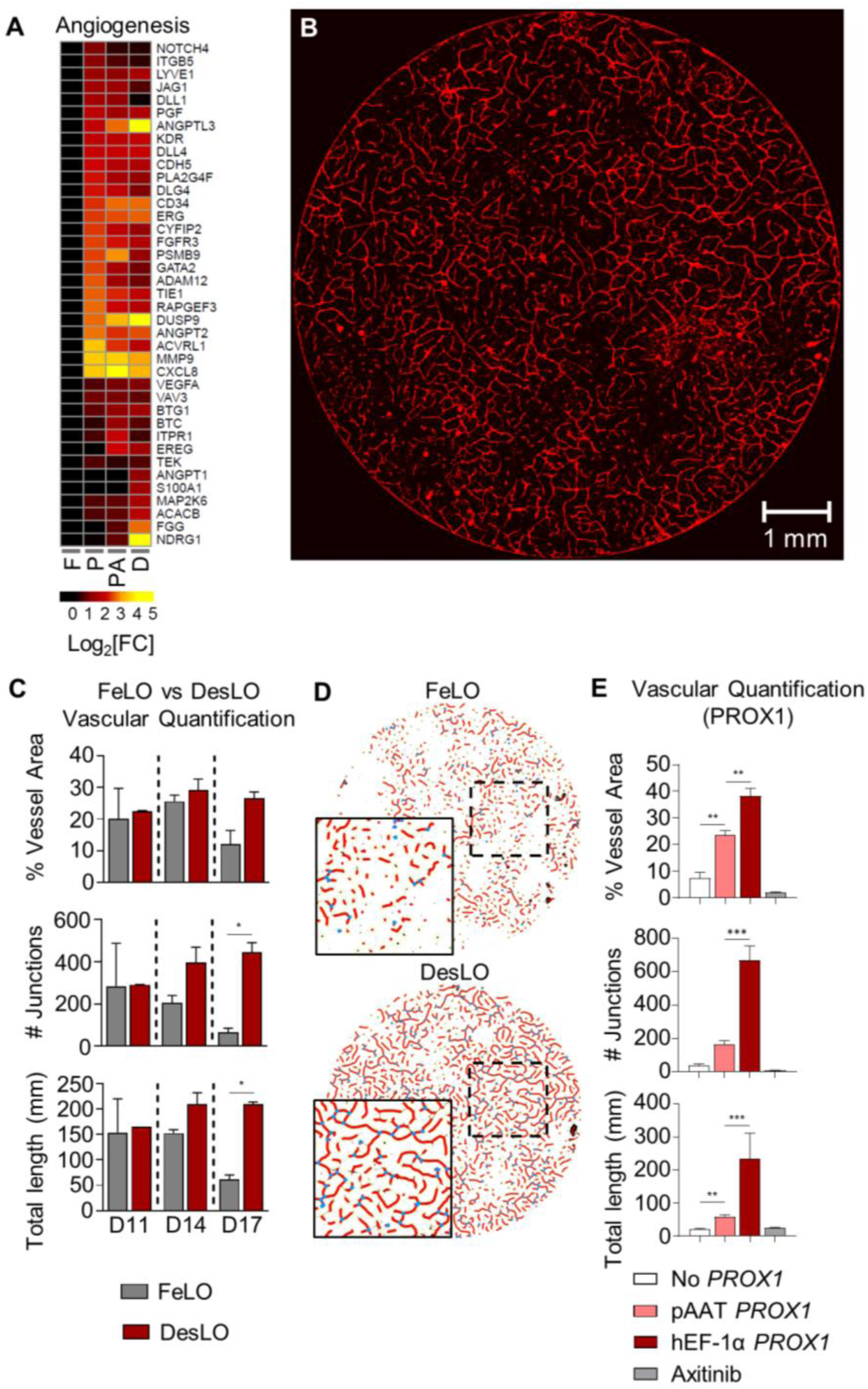
Vascular Development in DesLO. (A) Heatmap showing the expression relative to FeLO of angiogenesis-related genes in FeLO (F), PROX1 (P), PROX1+ATF5 (PA), and DesLO (D) transduced at day 5 (except FeLO) in day 17 tissues. (n=2, mean shown). (B) Immunofluorescence staining for CD31 showing vascular network on day 17 DesLO tissue. Image of 8mm diameter culture is shown. (C) AngioTool analysis of vessel area, vascular junctions, and total vessel length based on CD31 immunofluorescence staining of DesLO and control FeLO. Significance relative to control FeLO for each time point. D11, D14, D14: day 11, 14, 17 respectively. *p<0.05 (n=2). (D) Image interpretation of immunofluorescence staining of day 17 control FeLO and DesLO for CD31 generated by AngioTool analysis shows visual increase in vascular networks in DesLO. (E) AngioTool analysis of vessel area, vascular junctions, and total vessel length based on CD31 immunofluorescence staining of DesLO without *PROX1*, with *PROX1* driven by pAAT, and *PROX1* driven by hEF-1α with or without axitinib. **p<0.01, ***p<0.001 (n=3). Data are represented as mean ± SEM for D and E. See also Figure S6.

Given our findings on the key role of *PROX1* expression in induction of angiogenic cues, we asked whether confining *PROX1* upregulation to the hepatocyte population affected vascular formation. We generated DesLO as usual in parallel with tissues in which we either changed the *PROX1* promoter from constitutive hEF-1α to hepatocyte-specific pAAT or did not include a *PROX1* circuit at all. pAAT-*PROX1* DesLO showed a subtle but not significant decrease in *PROX1* expression relative to hEF-1α-*PROX1*, possibly due to fewer *PROX1*-expressing cells (**Figure S6C**). Compared to tissue without *PROX1*, pAAT-*PROX1* DesLO exhibited significantly improved vessel length, vessel area, and expression of *CD34* and *CD31*. However, the number of vessel junctions, vessel length, and vessel area were significantly compromised relative to DesLO with constitutively expressed *PROX1* (**Figures 6E, S6C, and S6D**). Therefore, these data suggest a key role for *PROX1* expression in both hepatocytes and non-hepatocyte cells in regulating angiogenesis of the developing liver.

To investigate the role of intercellular signaling in vascular formation we also treated hEF-1α *PROX1* DesLO cultures after day 7 with the small molecule axitinib, which selectively inhibits receptor tyrosine kinases (VEGFR1-3, PDGFRα,β and c-kit). Axitinib completely ablated formation of CD31^+^ cells and *CD34* and *CD31* expression (**Figures 6E, S6C, and S6D**), implicating such signaling as vital for the emergence of endothelium, which cannot be overcome by *PROX1* overexpression.

### Implantation in Mouse Models Provides a Therapeutic Advantage

To test the *in vivo* functionality of DesLO and its ability to engraft and provide therapeutic benefit, we implanted DesLO into mouse models of liver injury. We utilized FRGN mice (Wilson et al., 2014) in which liver injury is prevented with supplementation 2-(2-nitro-4-trifluoromethylbenzoyl)-1,3-cyclohexanedione (NTBC) in the drinking water (Vogel et al., 2004). Hepatocyte death and liver failure can be induced with removal of NTBC (**Figure 7A**) (see methods). Multiple DesLOs were placed over the mesentery vascular bed or packed beneath the renal capsule and secured using a fast-forming fibrin gel (**Figures 7B and S7A**).

**Figure 7.**
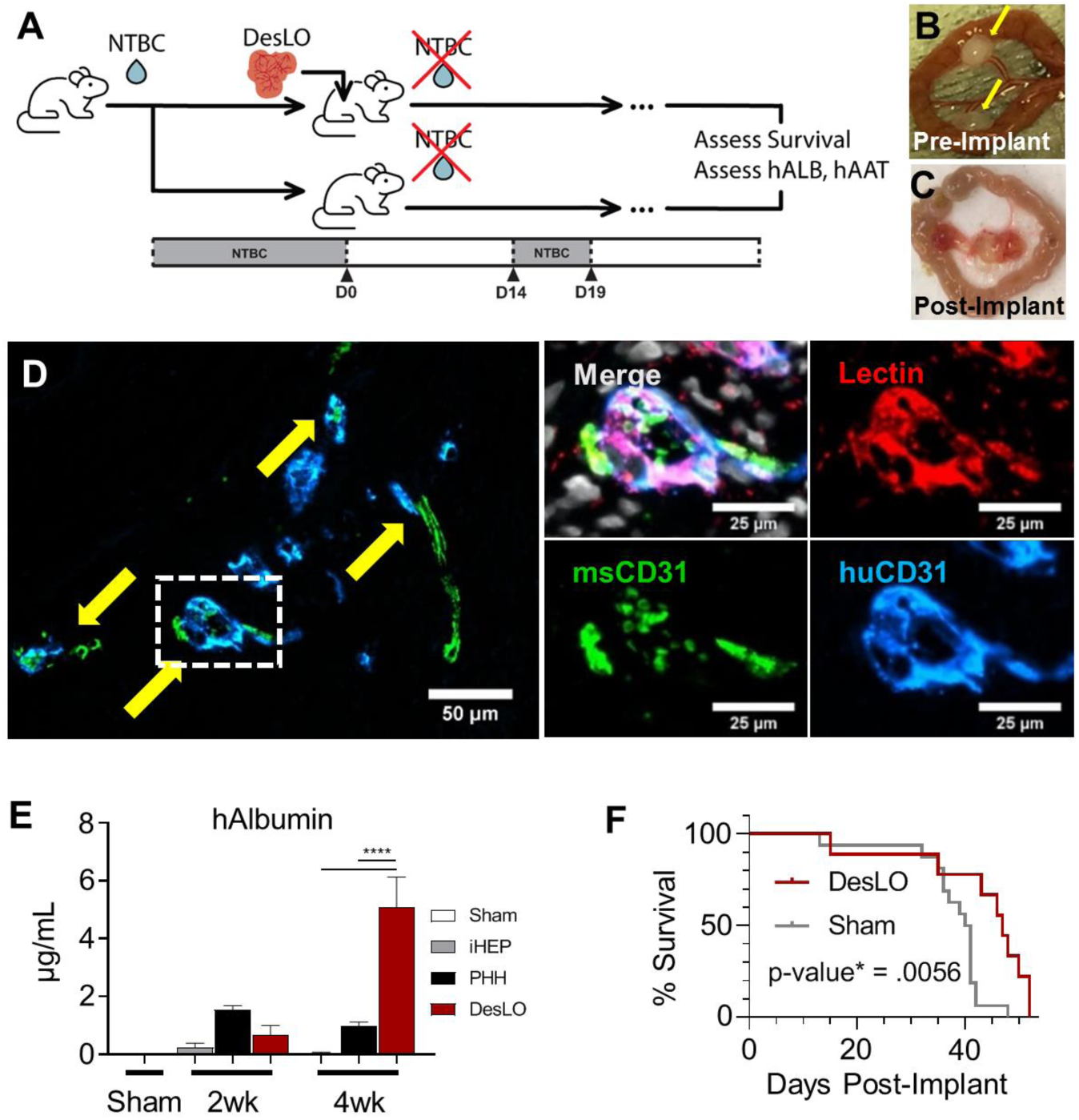
*In vivo* Function of DesLO after Implantation in Mice. (A) Schematic of implantation into FRGN liver injury mouse model *in vivo* experiments. (NTBC: nitisinone; hALB: human albumin; hAAT: human alpha-1 antitrypsin). (B) Fibrin-gelled DesLO tissue adhered to mesentery. Implants indicated by yellow arrows. (C) Mesentery-implanted DesLO harvested tissue after >4 weeks implanted shows tissue growth and vascularization. (D) Integration of human and mouse CD31 shown via immunofluorescence staining of paraffin section. (Left) Arrows indicate sites of overlap. (Right) Increased zoom of indicated selection denoted in left image by dotted rectangle. Lectin denotes vasculature from both species. (E) ELISA quantification of human albumin protein in mouse serum shows increase of human albumin between 2 and 4 weeks post-implantation of DesLO (n=5) at mesentery site, while detected human albumin decreases over this period in iHEP (n=7, from 3 lots) and PHH (n=6, from 3 lots). ****p<0.0001. (F) Kaplan Meier survival curve of FRGN mice with and without DesLO sub-renal implantations. Log rank p-value = 0.0056. Implant n=9, Sham n=11. Data are represented as mean ± SEM for E. See also Figure S7.

Harvested DesLO showed vascularization at the site of implantation (**Figures 7C and S7B**). Upon harvest of the implants, histology revealed human and mouse vascular integration via species-specific markers for CD31, with lectin staining the endothelial cells from both species (**Figure 7D**). Patches of hepatic cells producing human albumin were also identified from stained paraffin embedded sections of the recovered implant (**Figure S7C**). Secretion of hAlbumin and hAAT into host serum at two weeks post-implant was highest in the PHH control, but by four weeks post-implant PHH secretion had decreased while DesLO secretion increased to significantly higher levels (**Figures 7E and S7D**). This suggests that vascular integration may have facilitated efficient nutrient exchange and promoted implant growth and function. When we examined survival of mice receiving DesLO following complete removal of NTBC we found a moderate but statistically significant advantage over controls that was most evident after 40 days post liver injury (**Figure 7F**). Further optimization of the ectopic implantation procedure in terms of tissue mass, encapsulation methods, or implant site could facilitate improvements in survival outcome.

Implantation of tissues transduced with each factor alone revealed that hAlbumin secretion was largely driven by *PROX1* (**Figure S7E)**. hAlbumin levels were still substantially lower than those from DesLO implants however, suggesting that the components of the full circuit could synergize to confer *in vivo* functionality. To extend the applicability of these findings we used the TK-NOG mouse model and found similarly high levels of hAlbumin in the serum (**Figure S7F**). Taken together these data demonstrate the *in vivo* functionality and therapeutic potential of DesLO.

## DISCUSSION

In summary, we demonstrate advanced maturation and vascularization of human liver organoids via genetically guided engineering. We performed a comprehensive analysis of these organoids (DesLO) via multiscale assessment at the level of chromatin, evolved GRNs in single cells and tissue, as well as function *in vitro* and *in vivo*. In GRN meta-analysis, DesLO showed superior alignment to human liver and minimal aberrant signatures when compared with several other PSC-derived organoids. When tested *in vitro*, it performed at the level of primary adult hepatocytes for select metrics of function and exhibited a dynamic GRN state capable of response to environmental perturbations such as bile acid signaling. Strikingly, DesLO also showed improvement in vascular formation, another key objective in organoid engineering. Implantation *in vivo* demonstrated vascular integration, sustained secretion of human proteins, and therapeutic benefit in mice with liver damage. The ability to cryopreserve DesLO also indicates the feasibility of sharing experiments and resources across different laboratories. Despite these advances, DesLO could not match native hepatocytes in urea synthesis or CYP2C19 activity. These points emphasize the continuous need for future improvement using strategies such as iterative cycles of targeted genetic manipulation or combination with other approaches such as biomaterials, microfluidic devices, and growth factors. Yet, DesLO provides a proof of concept model for the feasibility of genomically-programmed formation and importantly maturation of complex PSC-derived tissues.

Maturation of some liver functions (e.g. CYP3A4 activity) continues months after birth, leaving us with a limited understanding of all *in vivo* microenvironmental cues necessary for faithful engineering of adult-like human livers. Our approach employed targeted activation of endogenous *CYP3A4* transcription in hepatocytes, which ignited further metabolic maturation in engineered DesLO, suggesting cell-intrinsic paths to override immature tissue states. Additionally, genetically engineering GRN state confers spatial control at the cellular level, which enabled confinement of CRISPR activation and assessment of *PROX1* function using pAAT (hepatocyte specific) in this study. Optimizing the CRISPR toolkit for potent modulation of endogenous loci is appealing in this context. Such methods circumvent any probable issues stemming from supraphysiological levels of overexpressed transcription factors, taking advantage of endogenous transcriptional programs and multiplexity (e.g. different gRNAs).

DesLO demonstrates vascular development without supplementation of angiogenic growth factors as a result of the genetically encoded molecular programs. This strategy reduces the need for complex media formulations that can support heterotypic cells (e.g. endothelial, pericyte, and epithelial), reducing the cost to produce and mature multicellular tissues. A similar approach was used recently via *ETV2*-expressing cells to vascularize cortical organoids (Cakir et al., 2019). Future work can take advantage of assembled layered genetic circuits (Kiani et al., 2014; Nissim et al., 2014) embedded in hiPSCs that can be activated based on cell state, decreasing the need for delivery of genetic cargo during differentiation in organoids. Such hiPSC lines with genetically embedded programs can also facilitate expansion and scaling of the derived organoids.

A rigorous, quantitative assessment of organoids is not frequently practiced in the field, making it difficult to perform side-by-side comparisons of organoids both with their *in vivo* counterparts and those generated through alternate protocols. Therefore, approaches that take into consideration tissue GRN relative to a compendium of native organs and other synthetic tissues can be beneficial for developing standardized methods to systematically evaluate *in vitro*-engineered organoids. To this end, the ultimate integration of comparative GRN assessment and modeling of tissue development with engineering technologies will offer a rational way to predictably steer *in vitro* organogenesis and establish assess-build-test cycles to improve the products of tissue engineering in the future.

## Author Contributions

M.R.E. conceived the initial idea and design of study and supervised the overall direction of the study. M.R.E, J.J.V., and R.L. designed the majority of experiments and were responsible for the interpretation with input from S.K.. R.L. and J.J.V. performed stem cell cultures, organoid engineering, and downstream assays. F.M., R.L., and J.J.V. produced genetic constructs with input from S.K.. J.J.V. and J.K. performed *in vivo* mouse studies. J.J.V. did the majority of single cell analyses with guidance and contributions from C.P., M.R.E., P.C., and Y.T.. Y.T. and P.C. performed CellNet analysis and helped with interpretation. D.C. and S.M.C.d.S.L. did the KeyGenes analyses and helped with interpretation. J.H. generated all schematic figures. S.L. provided SNP and ATAC-Seq analyses and helped with RNA-Seq analysis. J.J.V. contributed to ATAC-Seq analysis and together with R.L. participated in enrichr analyses. All authors contributed to the discussions during the study. M.R.E., R.L., J.J.V. and S.K. wrote the manuscript with edits and comments from J.H., F.M., P.C., Y.T., D.C., S.M.C.d.S.L., S.L., and C.P..

## Acknowledgements

We thank all members of Ebrahimkhani and Kiani laboratories for discussions and assistance during this study. We also thank the Mayo Clinic Histology Core Laboratory for assistance with embedding and sectioning of tissue samples as well as the ASU Department of Animal Care and Technologies for assistance in mouse husbandry. We also thank the UCLA Technology Center for Genomics and Bioinformatics, Novogene, and Genewiz for help with RNA sequencing. This work was supported by an R01 from National Institute of Biomedical Imaging and Bioengineering (EB028532), an R01 from the Narional Heart, Lung, and Blood Institute (HL141805) and the New Investigator Award from Arizona Biomedical Research Council (ADHS16-162402) to M.R.E. as well as support from the Pittsburgh Liver Research Center (NIH-NIDDK P30DK120531). P.C. was supported by R35GM124725.

## Lead contact and materials availability

RNA-Seq, scRNA-Seq, and ATAC-Seq data is currently being prepared for submission to repository and will be available. All data is available upon request. Information and requests for resources and reagents should be directed to and will be fulfilled by the Lead Contact, Mo R. Ebrahimkhani

## Conflict of interest

M.R.E, S.K., P.C., J.J.V., and R.L. have submitted a patent (WO2019237124) for the work included in this publication.

---

**Figure S1.**
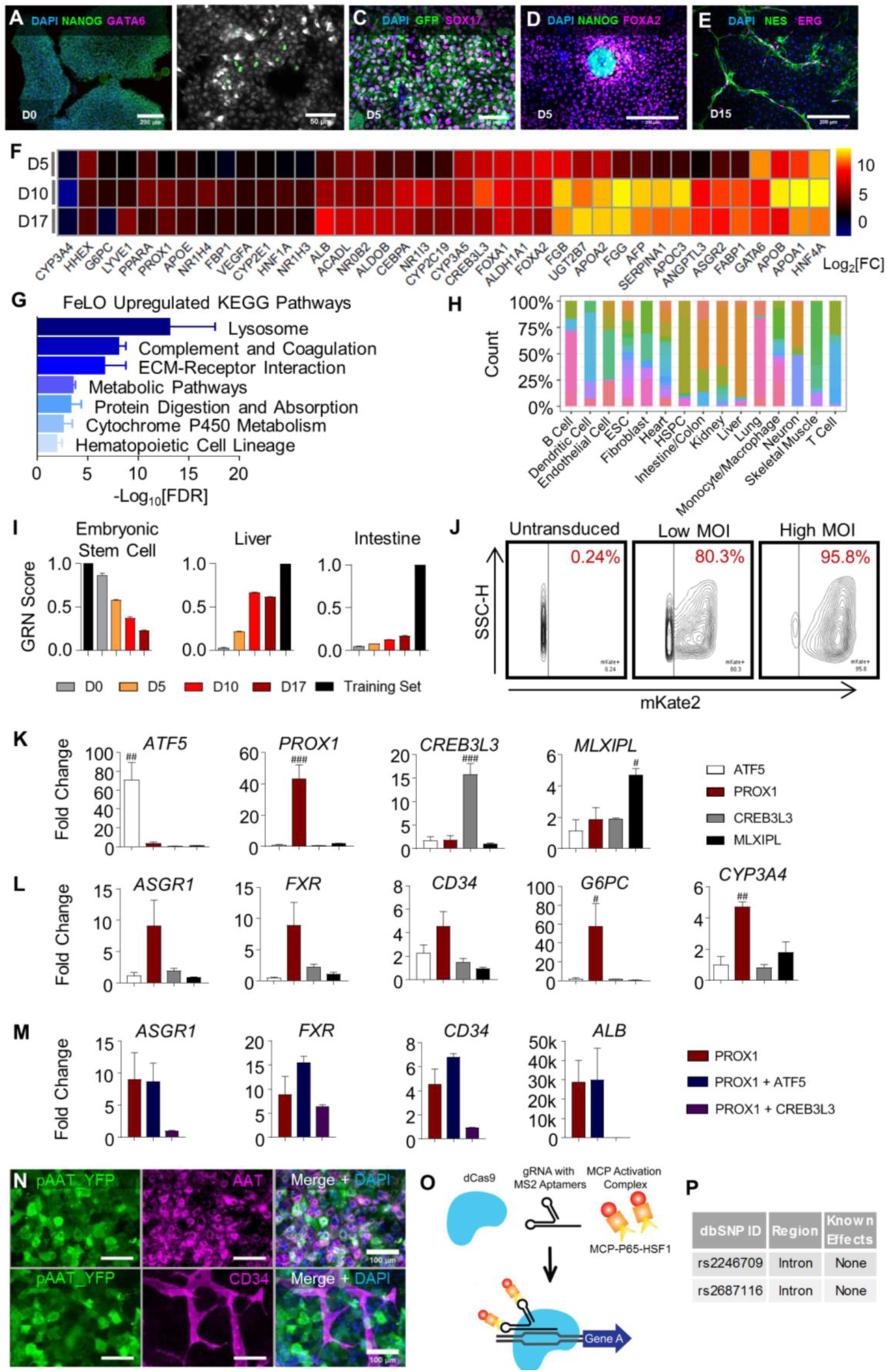
Generation and Assessment of Fetal Liver Organoid (FeLO). Related to Figure 1. (A-E) Immunofluorescence stains of pluripotency marker NANOG and GATA6 at day 0 shows no expression of GATA6 prior to induction with dox (A). Mesendodermal markers appear during dox-induced differentiation at day 1 (B) and day 5 (C). NANOG and FOXA2 are mutually exclusive (D). NES+ pericyte and ERG+ endothelial cells associate with each other (E). (F) Heatmap showing increased expression of hepatic developmental and functional genes on days 5, 10, and 17. Expression is represented as log2 of the fold change (log2[FC]) over non-induced cultures. (n=2, mean shown). (G) Enrichr-generated -log10[FDR] for day 17 FeLO KEGG pathways upregulated over hiPSC. (n=2). (H) Distribution of studies contributing to each cell/tissue type in the CellNet training data. The y-axis represents the proportion of samples of the cell type/tissue indicated on the x-axis that come from each of the 97 studies, as indicated by the color. (I) Graphical depiction of the CellNet GRN scores relative to the embryonic stem cell, liver, and intestine training sets. *Training set refers to the library of the specified cell type/organ. D0, D5, D10, and D17: day 0, 5, 10, and 17 respectively. (n=2). (J) FACS scatter plots showing transduction efficiency by mKate2 expression in untransduced, low MOI (9 MOI), and high MOI (150) conditions. (K) qPCR data for transcription factor screening in FeLO validating upregulation of each individual factor. Reference is untransduced. # = p<0.05, ## = p<0.01, ### = p<0.001 compared with each other condition (n=3). (L) qPCR data for transcription factor screening in FeLO of increase in *ASGR1, FXR, CD34, G6PC*, and *CYP3A4* in ATF5, PROX1, CREB3L3, or MLXIPL engineered tissues. Reference is untransduced. # = p<0.05, ## = p<0.001 over each other condition (n=3). For PROX1 engineered group, the same data are shown in Figures 1J, 1K, and S1M for the indicated targets for the purposes of clear comparison. (M) qPCR data for transcription factor screening in FeLO of PROX1 and PROX1 with ATF5 or CREB3L3 co-expression for *ASGR1, FXR, CD34*, and *ALB*. Reference is untransduced. (n=3). *ALB* data for PROX1 engineered group is the same as that shown in Figure 1I for the purpose of clear comparison. (N) Immunofluorescence staining of AAT and YFP in pAAT YFP-transduced *GATA6*-engineered hiPSC cultures shows that AAT promoter-driven YFP co-expresses with AAT and is not expressed in CD34+ endothelial cells. (O) Schematic of CRISPRa approach for building SynTF(*CYP3A4*) using dCas9 with gRNA containing MS2 aptamers and MCP fused to transcriptional activators (MCP-P65-HSF1) to upregulate target genes. (P) Table with information regarding the SNPs in *CYP3A4* in our cell line. Data are represented as mean ± SEM for G, I, K-M.

**Figure S2.**
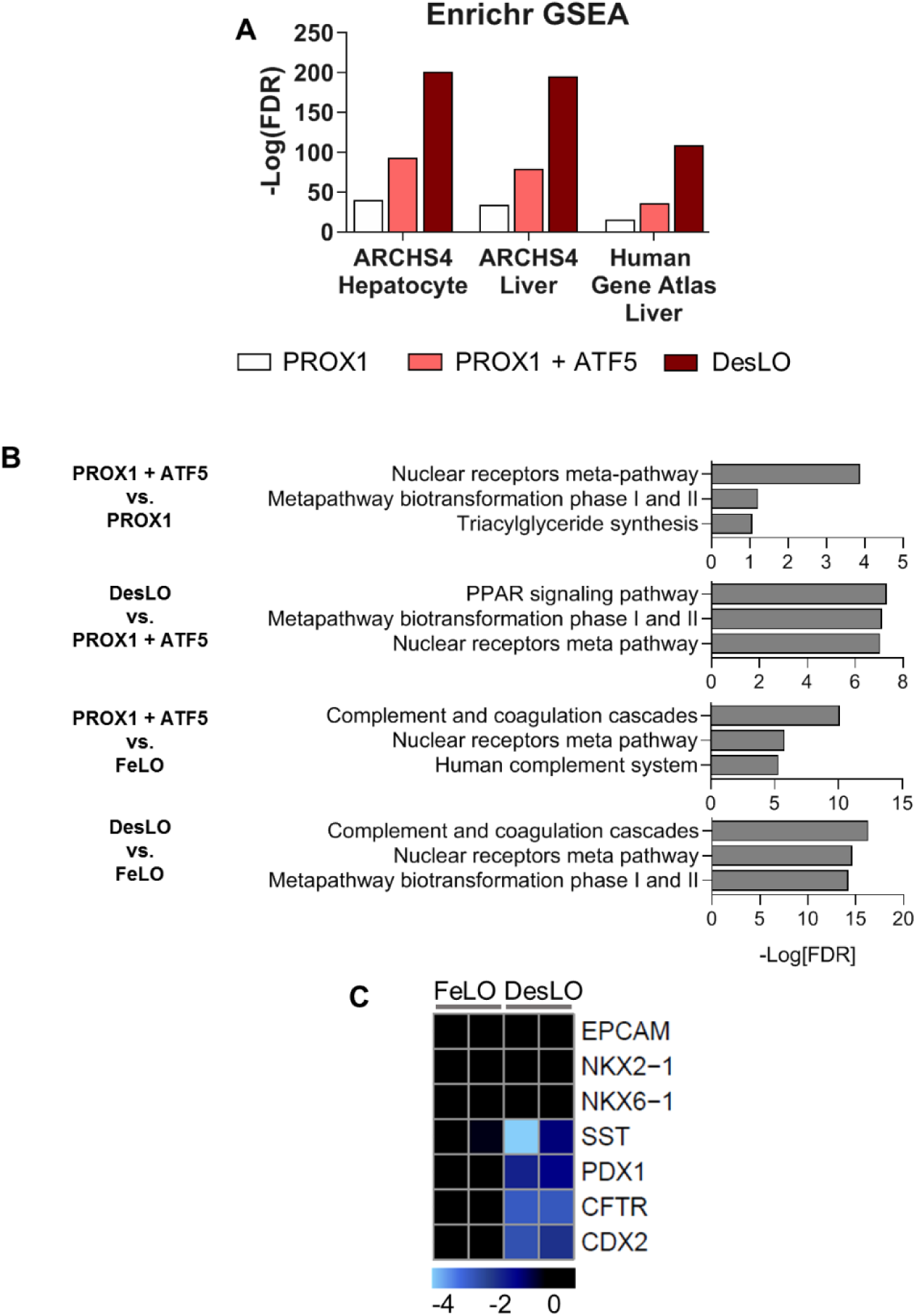
Genetic Engineering Improves Hepatic Identity and Tissue Maturation. Related to Figure 2. (A) Log10[FDR] of day 17 upregulated gene (>2 fold-change over age-matched FeLO) alignment with the ARCHS4 and Human Gene Atlas transcriptome libraries for hepatocyte and liver when transducing different gene circuits. (B) Log10[FDR] for enriched pathways at day 17 for the given comparisons when transducing different gene circuits using the WIkiPathways 2019 library and Enrichr. (C) Heatmap of gene expression in FeLO and DesLO from RNA-Seq data highlighting decreased DesLO expression of aberrant lineages (values are relative to FeLO).

**Figure S3.**
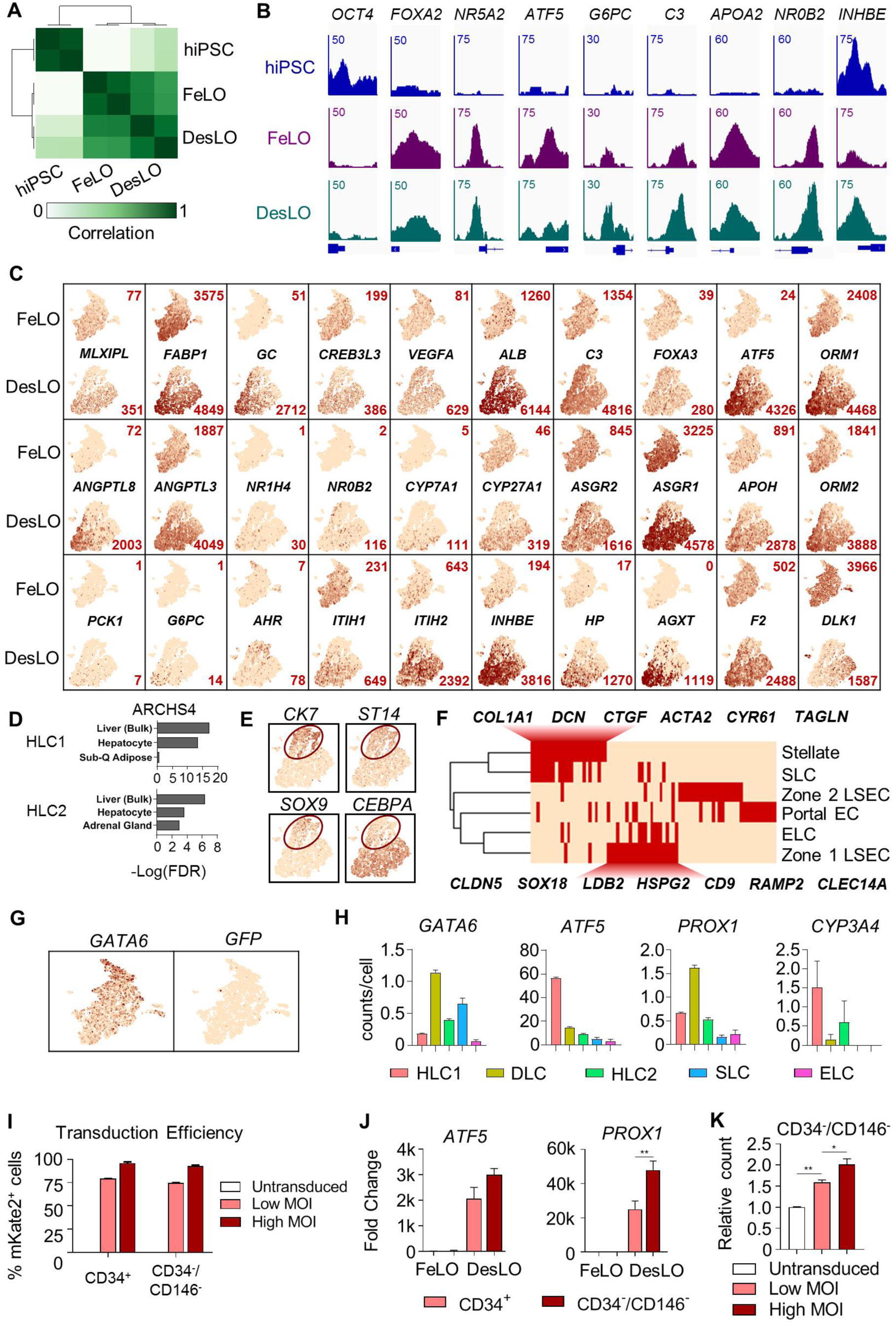
Cross-study Comparison of *in vitro*-derived Organoids, ATAC-Seq, and scRNA-Seq Analysis of DesLO. Related to Figure 3. (A) Correlation plot of ATAC-Seq data for hiPSCs, FeLO, and DesLO samples. (B) ATAC-Seq results showing changes in chromatin accessibility in *OCT4, FOXA2, NR5A2, G6PC, C3, APOA2, NR0B2*, and *INHBE* promoter regions in hiPSCs, FeLO, and DesLO. The scale for the y-axis is shown for each graph in reads. (C) tSNE plots of FeLO and DesLO for notable liver genes. The numbers indicate the number of cells in the data containing >1 UMI count of the indicated liver genes. Color scale is set per gene to portray FeLO and DesLO comparison. (D) Enrichr data showing the -log10[FDR] associated with the HLC1 and HLC2 clusters against the ARCHS4 tissue library (qualified by >1.6 fold change over average expression of all other clusters). (E) tSNE plots showing biliary/ductal cell gene expression of *CK7*+/*ST14*+/*SOX9*+/*CEBPA*-. (F) Binary heat map for the indicated tissues and tissue-specific genes shows that day 17 DesLO clusters align with the putative cell identities. A set of key overlapping genes are displayed above (SLC-stellate overlap) and below (ELC-zone 1 EC overlap) the heatmap. Selection and expression criteria explained in materials and methods. (G) tSNE plots of day 17 FeLO show endogenous *GATA6* expression with no remaining *GATA6-2A-EGFP* transgene expression shown by the absence of GFP signal. (H) Mean UMI counts of DesLO per cell by cluster of *GATA6, ATF5, PROX1*, and *CYP3A4*. (n = number of cells per cluster, shown in Figure 3H). (I) FACS data for transduction efficiency by MOI for the cell populations shown. Details are listed in the Methods section. (n=3 except n=2 for High MOI). (J) qPCR data for *ATF5* and *PROX1* showing differential transgene genomic integration rates among the cell populations shown. Y axis is relative expression against CD34-/CD146-FeLO. **p<0.01 (n=3). (K) FACS data showing relative proportion of CD34-/CD146-hepatic population in untransduced, low MOI (9), and high MOI (150). *p<0.05, **p<0.01. (n=3 except n=2 for High MOI). Data are represented as mean ± SEM for H-K.

**Figure S4.**
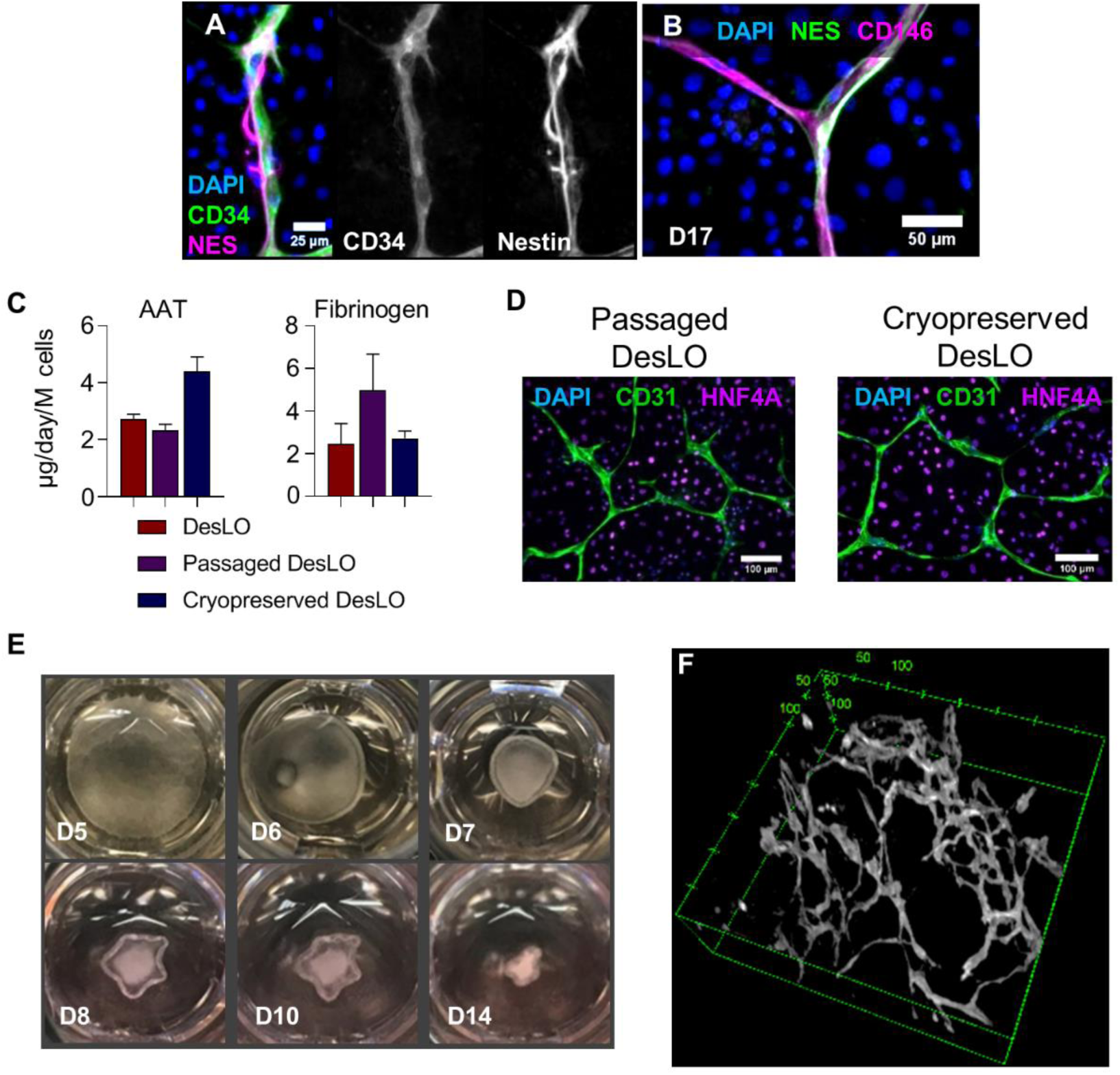
Immunofluorescence and Functional Comparison of DesLO, iHEP, and PHH cultures. Related to Figure 4. (A-B) Immunofluorescence staining of day 17 DesLO cultures highlighting the close association of CD34+ and NES+ cells (A) and the co-expression of NES and CD146 (B) (D17: day 17). (C) AAT and Fibrinogen ELISA measurements of unpassaged, passaged, and cryopreserved DesLO cultures (n=3). (D) Immunofluorescence images showing intact vasculature of unpassaged, passaged, and cryopreserved DesLO cultures. (E) Time lapse of DesLO contraction after day 5 transduction and seeding on Matrigel gels. (F) 3-D rendering of CD31-stained vasculature of gel surface-generated day 17 DesLO tissue. Scale units are in microns. Data are represented as mean ± SEM for C.

**Figure S5.**
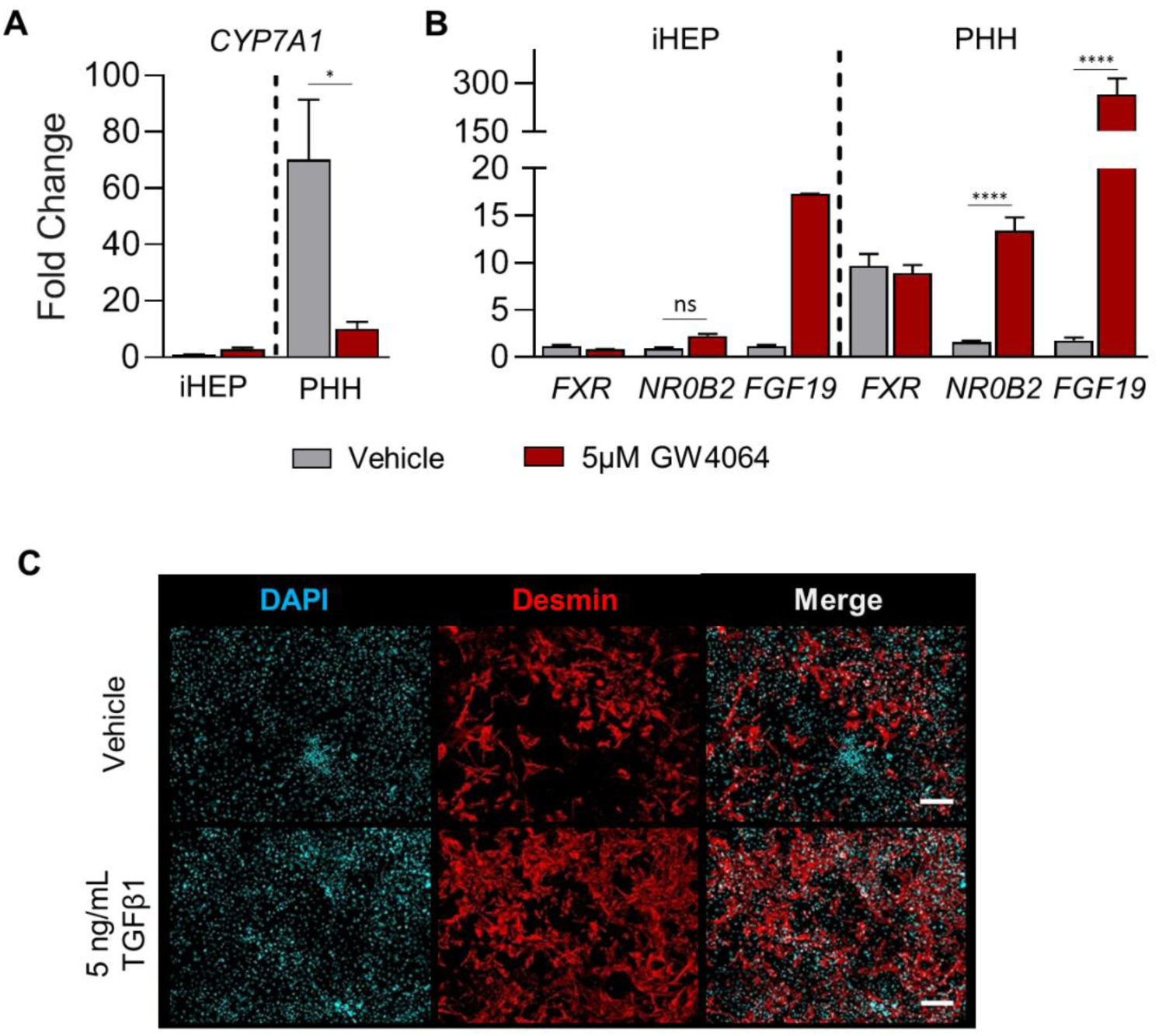
Modeling FXR Signaling and Stellate Cell-like Function in DesLO. Related to Figure 5. (A) qPCR showing fold change over iHEP vehicle control of *CYP7A1* in iHEP and PHH with addition of GW4064. *p<0.05 (n=4 for PHH from 2 lots and n=2 for iHEP). (B) qPCR showing fold change over iHEP vehicle control of *FXR, NR0B2, FGF19* in iHEP and PHH with addition of GW4064. ****p<0.0001, ns: not significant (n=4 for PHH from 2 lots and n=2 for iHEP). (C) Immunofluorescence staining for Desmin in DesLO after supplementing 5ng/mL TGFβ1 for 3 days. Data are represented as mean ± SEM for A and B.

**Figure S6.**
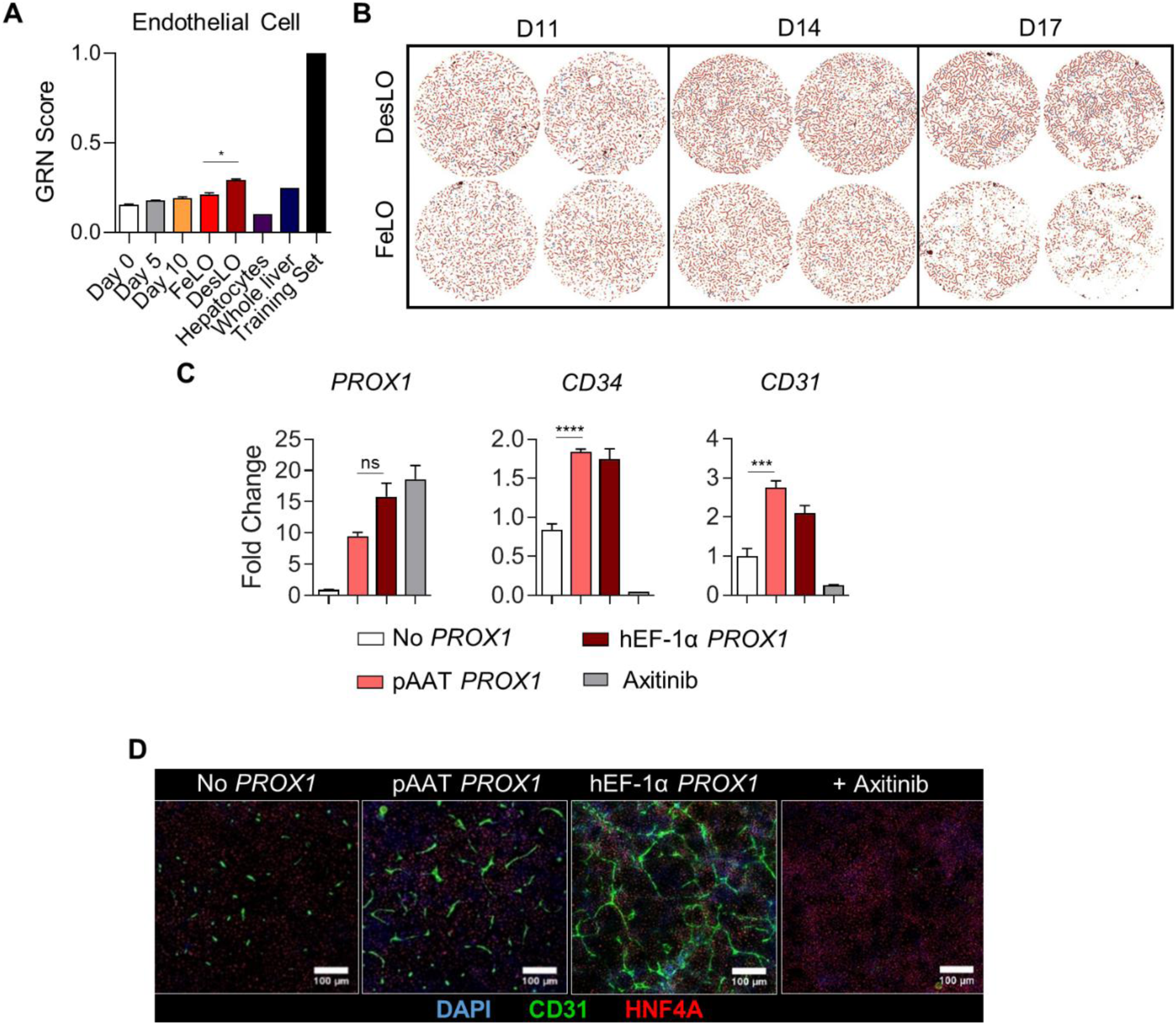
Vascular Development in DesLO. Related to Figure 6. (A) CellNet GRN scores for FeLO timelapse samples, DesLO, freshly isolated primary human hepatocytes, and whole liver tissue relative to training set. Day 0 is uninduced hiPSC and Day 5 and Day 10 are FeLO at each time point. FeLO and DesLO samples are day 17. Training set represents human endothelial cells. *p<0.05, two tailed t test (n=2). (B) Binary image interpretation of each CD31 staining sample from of day 11 (D11), day 14 (D14), and day 17 (D17) of control FeLO and DesLO generated by AngioTool analysis to determine vascular quantification metrics. (C) qPCR data for *PROX1, CD34*, and *CD3*1 expression in DesLO without *PROX1*, with *PROX1* driven by pAAT, and *PROX1* driven by hEF-1α with or without axitinib. ***p<0.001, ****p<0.0001, ns = not significant (n=3). (D) Immunofluorescence staining of CD31 and HNF4A in DesLO with no *PROX1*, pAAT *PROX1*, hEF-1α *PROX1*, and hEF-1α *PROX1* following axitinib treatment in DesLO with hEF-1α *PROX1*. Data are represented as mean ± SEM for A and C.

**Figure S7.**
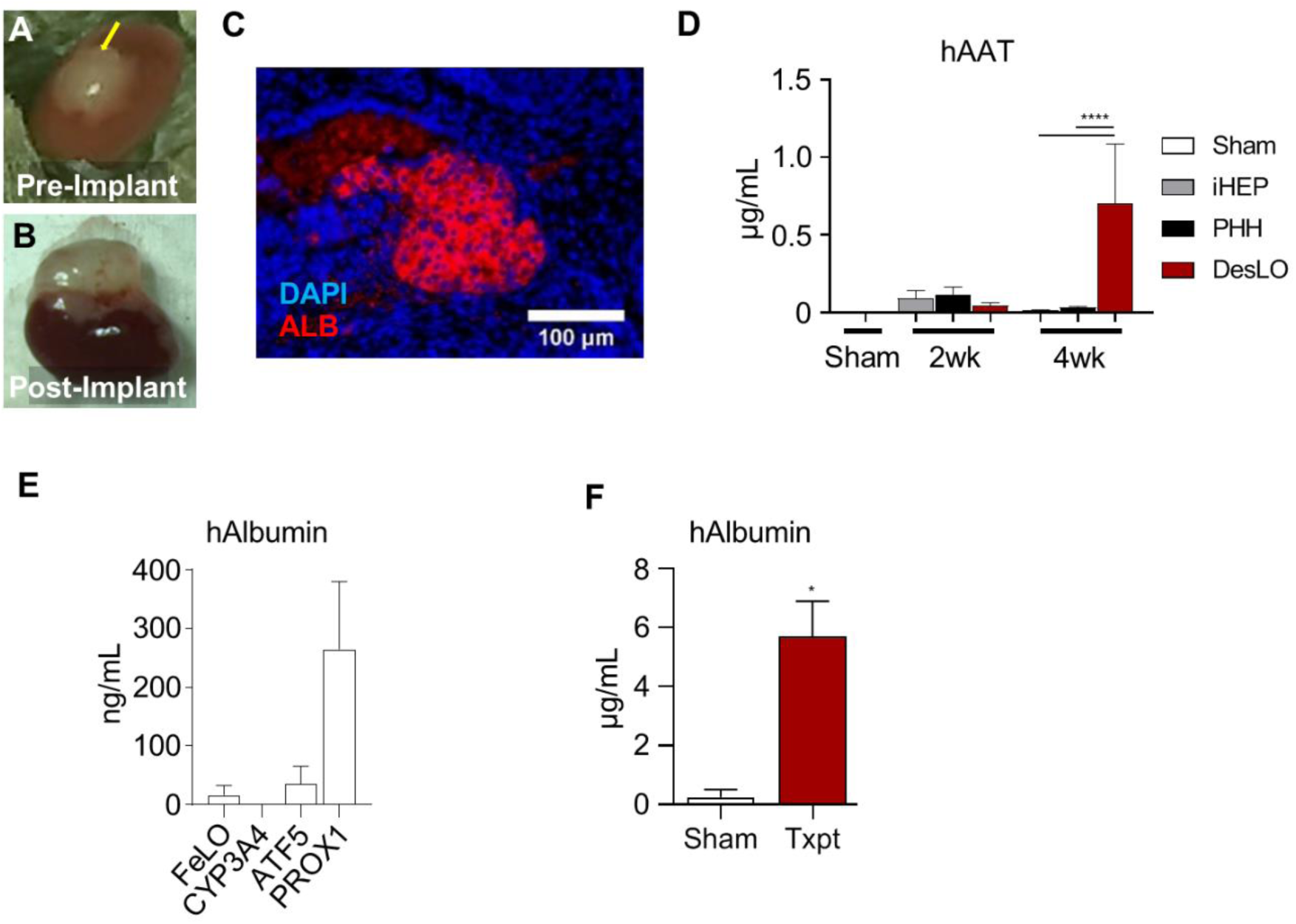
*In vivo* Function of DesLO after Implantation in Mice. Related to Figure 7. (A) Fibrin-gelled DesLO tissue inserted beneath renal capsule. Implant indicated by yellow arrow. (B) Renal capsule-implanted DesLO tissue growth and vascularization after 4 weeks of implantation in FRGN liver injury mouse model. (C) Human albumin staining of paraffin embedded tissue section from harvested renal capsule implant. (D) ELISA quantification of human AAT protein in mouse serum shows increase of human AAT between 2 and 4 weeks post-implantation of DesLO at mesentery site, while detected human AAT decreases over this period in all other conditions. ****p<0.0001, iHEP (n=7, from 3 lots) and PHH (n=6, from 3 lots). (E) ELISA quantification of human albumin in mouse serum shows that tissue transduced with *PROX1* generates more albumin than FeLO and tissue transduced with *ATF5* or SynTF(*CYP3A4*) after 4 weeks of implantation. (n=3-4 except n=2 for *ATF5*-transduced DesLO). (F) Human albumin measurements in mouse serum of TK-NOG liver injury model between 35-45 days post-implant (blood taken on day of euthanization). *p<0.05, two tailed t-test (n=3). Data are represented as mean ± SEM for D-F.

