## Supplementary Material for "Synthetic Maturation of Multilineage Human Liver Organoids via Genetically Guided Engineering"

### Supplement includes the following:

- EXPERIMENTAL MODEL AND SUBJECT DETAILS
- METHOD DETAILS
- QUANTIFICATION AND STATISTICAL ANALYSIS
- KEY RESOURCE TABLE

### EXPERIMENTAL MODEL AND SUBJECT DETAILS

#### Mice

All animal husbandry and experiments were performed after ethical committee review and in accordance to The Institutional Animal Care and Use Committee at Arizona State University. The *Fah$^{-/-}$ / Rag1$^{-/-}$ / Il2rgnull* mouse strain on NOD background (FRGN) was purchased from Yecuris. The knockout of Fah necessitates the constant supplementation of 2-[2-nitro-4-(trifluoromethyl)benzoyl]cyclohexane-1,3-dione (NTBC, or nitisinone, 16 mg/L) in the feeding water in order to prevent accumulation of toxin in the liver. Both and female FRGN mice age were used for this study (with studies starting at 8-12 weeks of age), no significant differences were detected between male and female results. The NOD.Cg-*Prkdc*$^{scid}$*Il2rg*$^{tm1Sug}$ Tg(Alb-TK)7-2/ShiJic (TK-NOG) strain was purchased from Taconic. TK-NOG mice have a hepatocyte specific expression of a Herpes Simplex Virus-1 Thymidine Kinase (TK) downstream of the albumin promoter which, upon systemic delivery of Ganciclovir (GCV), exhibits hepatocyte-specific ablation. Mice were bred (FRGN mice only, TK-NOGs from Taconic cannot be bred) housed, fed, and monitored in accordance with the protocols approved by the Institutional Animal Care and Use Committee at Arizona State University and all animals were carefully monitored for signs of morbidity and discomfort by research and veterinary staff. Experiments utilized Male only for TK-NOG and both male and female for FRGN. There were no differences noticed in results based on sex.

### METHOD DETAILS

#### Lentiviral Production and Titration

HEK293FT cells (Life Technologies) were grown according to the manufacturer's instructions in a humidified incubator at 37 °C with 5% $CO_2$. The day before transfection 8 million HEK293FT cells were seeded on a collagen I-coated (Gibco A10483-01) 15 cm$^2$ tissue culture treated cell culture dish. On the day of transfection cells were co-transfected with 15 µg psPAX2 (Addgene Plasmid 12260), 3.75 µg pCMV-VSV-G (Addgene Plasmid 8454), and 11.25 µg of the plasmid to be packaged using 90 µg of linear polyethylenimine (Polysciences, Inc 23966-1). Medium was changed the next morning and the supernatant was collected after 48 and 72 hours. Pooled supernatant was filtered through a 0.45 µm low protein binding filter (Corning) and concentrated in an Amicon Ultra 15 filter columns (100 kDa cutoff, Millipore) at 4,000*g* for 23 minutes. The concentrated virus was then aliquoted, snap frozen, and stored at −80 °C. Lentiviral concentrate was diluted 2000-fold in phosphate buffered saline (PBS) and titered via qRT-PCR using a commercially available kit (ABM LV900). Titers were calculated according to manufacturer's instructions.

#### Vector Design and Construction

*sgRNA, MPH, and dCas9 Lentiviral Vectors*

The MS2-P65-HSF1-GFP (Addgene plasmid ID: 61423) MCP-fused transcriptional activator (MPH) was amplified and sub-cloned into a gateway entry vector via golden gate-based reaction for further cloning into a lentiviral gateway destination vector. To generate U6-sgRNA-MS2 plasmids, 20bp guide sequences were inserted into the sgRNA-MS2 cloning backbone (Addgene plasmid ID: 61424) at the BbsI site via golden gate-based reaction. After screening, U6-sgRNA-MS2 for two target sites were amplified and sub-cloned into a gateway entry vector via golden gate-based reaction. The U6-sgRNA-MS2 (x2) and MPH entry vectors were cloned into a lentiviral destination vector via gateway cloning. dCas9 (Addgene plasmid ID: 47319) was amplified and sub-cloned into a gateway entry vector via golden gate-based reaction and then cloned with the gateway entry vector for the AAT promoter into a lentiviral gateway destination vector using gateway cloning.

*Transcription Factor Vectors*

*ATF5* transcript variant 1 cDNA was purchased from Origene (Cat. #RC200081), *PROX1* transcript variant 2 cDNA was purchased from GeneCopoeia (Cat. #F0925), *CREB3L3* complete CDS cDNA was purchased from DNASU (Clone ID# HsCD00080068), and *MLXIPL* transcript variant 1 cDNA was purchased from DNASU (Clone ID# HsCD00820703). These plasmids were individually amplified from their respective vectors and sub-cloned into gateway entry vectors via golden gate reaction. The *PROX1* entry vector was cloned with gateway entry vectors for either the hEF-1α or AAT promoter into a lentiviral gateway destination vector via gateway cloning. The *ATF5*, *CREB3L3*, and *MLXIPL* entry vectors were cloned with a gateway entry vector for the hEF-1α promoter into a lentiviral gateway destination vector via gateway cloning. For constructing the dox-inducible expressing *GATA6* vector, pENTR_L1_hGATA6-2A-EGFP_L2 previously published (Guye et al., 2016) was cloned into an All-in-One PiggyBac transposon destination vector from Addgene (Addgene plasmid ID: 80479) via gateway reaction.

**Guide RNA Design**

Guide RNA (gRNA) was designed to target the proximal promoter region of *CYP3A4*. Target loci were selected using the SAM gRNA design tool (Konermann et al., 2014) or custom designed.

**Magnetic Depletion of TRA-1-60 Expressing Cells**

TRA-1-60-expressing cells were depleted from cultures using the Miltenyi MACS separation system after 5 days of doxycycline induction. Cells were dissociated with accutase and depletion was performed using TRA-1-60 antibody-conjugated magnetic beads (Miltenyi Biotech) according to the manufacturer's instructions.

**Lentiviral Transduction**

Pre-aliquoted lentiviral concentrate was thawed quickly and maintained on ice before use. hiPSCs were transduced as single-cell suspensions in Matrigel-coated 48-well cell culture plates in mTeSR-1 containing 8μg/mL polybrene,10 μM Y-27632, and 1μg/mL doxycycline. The medium was changed the following day. Multiplicity of infection (MOI) was determined by dividing the number of lentiviral infectious units (calculated from the lentivirus titration step) by the number of cells being seeded with the virus. Where indicated, the lentiviral MOI was 9 for low MOI and 150 for high MOI.

**Cell Culture**

All cells and tissues were cultured in a humidified incubator at 37 °C and 5% $CO_2$. Our hiPSC lines were cultivated under sterile conditions in mTeSR-1 (Stem Cell Technologies, Vancouver) changed daily. Tissue culture plates were coated for 1 hour at room temperature with BD ES-qualified Matrigel (BD Biosciences) diluted according to the manufacturer's instructions in ice cold DMEM/F-12 with 15 mM HEPES medium (Thermo Scientific). Routine passaging was performed by incubating hiPSC colonies for 5 minutes in Accutase (Sigma) at 37 °C, collecting the suspension and adding 5mL DMEM/F-12 medium containing 10 μM Y-27632, centrifuging at 300*g* for 5 minutes, and resuspending in DMEM/F-12 supplemented with 10 μM Y-27632 for counting. Cells were seeded at a cell density of 25,000 cells per $cm^2$.

Primary cryopreserved human hepatocytes were obtained from Thermo Fisher, Lonza, Zenbio, and the MGH Cell Resource Core (refer to Key Resources Table for catalog and lot information). Hepatocytes were thawed, seeded, and cultured according to distributors' instructions with the following held constant between both distributors: hepatocytes were seeded in rat tail collagen I (Thermo scientific)-coated 48-well plates at a seeding density of 175,000 viable hepatocytes per $cm^2$, human cryopreserved hepatocyte thawing medium (Lonza) was used for recovery following thaw, Thawing/Plating Cocktail A + 5% fetal bovine serum was used for seeding, and Cell Maintenance Cocktail B was used for maintenance of hepatocytes with William's Medium E (Thermo Scientific) as basal medium. Function was assessed via staining as well as ELISA, CYP3A4 activity, and urea production by day 4 of culture using at least 3 unique donors with age and sex matching when possible. One lot was used for CYP2C19 assessment. Results were normalized by number of seeded cells. Human stem cell-derived hepatocytes (iHEP) were purchased from Takara Bio Inc. (2 lots) and Stem Cell Technologies (1 lot), thawed and cultured according to the manufacturer's protocols, and assessed at 8 days. HEK293FT cells (Thermo Scientific) were cultured under sterile conditions according to the user guide instructions.

#### *GATA6*-Engineered Cell Line Generation

rtTA expressing PGP1 hiPSCs previously generated (Guye et al., 2016) were transfected using Lipofectamine 3000 (ThermoFisher Scientific) with Super PiggyBac Transposase (System Biosciences) and the PiggyBac transposon vector with hGATA6-2A-EGFP under control of the tetracycline responsive element promoter. Transfected cells were selected by adding 0.5μg/mL puromycin to the mTeSR1 maintenance medium.

#### Generation of FeLO and DesLO

FeLO is produced as explained before (Guye et al, 2016) except we used the engineered cells developed by *GATA6* transposition. The *GATA6*-engineered hiPSCs were seeded at a density of 25,000 cells per $cm^2$ in mTeSR-1 supplemented with 10 μM Y-27632. The next day, the medium was changed to mTeSR-1 with 1 μg/mL doxycycline to induce expression of the *GATA6* transgene and replaced daily for 5 days. On day 5 the medium was changed to APEL, fully defined, serum-free and animal component-free medium (Stem Cell Technologies, Vancouver), and replaced daily until the day of harvest. For DesLO generation, cells were dissociated with accutase, depleted of TRA-1-$60^+$ undifferentiated cells, transduced as described above with lentiviruses (*PROX1*/*ATF5* transgenes and SynTF(*CYP3A4*)) on day 5, and seeded at a density of 300,000 cells per $cm^2$ in mTeSR-1 supplemented with 1μg/mL doxycycline and 10 μM Y-27632. The next day, medium was replaced with mTeSR-1 with 1 μg/mL doxycycline. The medium was switched the following day to APEL and replaced daily. FeLO controls were generated through the same protocol as DesLO described above but were either not transduced or transduced with dCas9 (without gRNA) and mKate2 lentivirus. Samples were harvested at day 17 of culture unless otherwise indicated. For experiments in which axitinib (Cell Signaling Technology Cat. #12961S) was used, it was administered in the culture medium upon the switch to APEL on day 7 at 50 nM. For FXR regulation assessment studies day 14 DesLO media was supplemented with either 1:1000 dimethylsulfoxide (DMSO) vehicle, 5 μM GW4064, 50 μM Chenodeoxycholic acid (CDCA), or 25 μM FGF19 with media refreshed daily until harvest at day 17. For fibrosis induction and mitigation experiment, day 14 DesLO media was supplemented with either 1:1000 dimethylsulfoxide (DMSO) vehicle, 5 ng/mL TGFβ1, or 5 ng/mL TGFβ1 and 5 μM GW4064 together with media refreshed daily until harvest at day 17.

#### DesLO Passaging

DesLO was generated as described, but rather than switching to APEL on day 7 of culture, tissue was maintained on mTeSR-1 supplemented with with 1μg/mL doxycycline until day 10. On day 10 tissue was dissociated with accutase and reseeded at 300,000 cells per $cm^2$ in mTeSR-1 supplemented with 1μg/mL doxycycline and 10 μM Y-27632. From this point the DesLO were cultured as previously described post-transduction for twelve days with APEL.

#### DesLO Freezing and Thawing

DesLO was generated as described, but rather than switching to APEL on day 7 of culture, tissue was dissociated using accutase and frozen in mFresR cryopreservation medium supplemented with 10 μM Y-27632. Cells were thawed and seeded into the same size well from which the vial was frozen. The next day, culture medium was switched to APEL and tissue was cultured for nine more days until day 17 of total culture.

**MACS Bead Isolation of FeLO and DesLO**

On day 15 of culture FeLO and DesLO were dissociated using trypsin. MACS bead isolation for $CD34^+$ cells was performed according to manufacturer's instructions (Miltenyi 130-100-453). MACS bead isolation for $CD146^+$ cells was performed on the $CD34^-$ population according to manufacturer's instructions (Miltenyi 130-093-596). DNA for the $CD34^+$ the $CD34^-/CD146^-$ populations was extracted using the Allprep DNA/RNA Plus Micro Kit (QIAGEN, Cat. #80284). Quantitative PCR was performed on genomic DNA with expression normalized to genomic Albumin and relative gene expression calculated using $2^{-\Delta\Delta CT}$ method.

**Flow Cytometry**

To test transduction efficiency DesLO was generated as described, with the exception that also included was an expression vector for mKate2 under control of the hEF-1α promoter. Transduced (DesLO) and untransduced (FeLO) tissues were dissociated at day 15 of culture using trypsin (Gibco 15400054) and analyzed using a Thermo Fisher Attune NxT Cell Analyzer flow cytometer. Analysis were performed using FlowJo 2.

**Tissue Harvest, RNA Extraction, qRT-PCR**

Tissues were harvested and lysed by adding 500μL Trizol (Life Technologies) directly to the tissue culture well and storing at -80°C. For extraction, lysate was thawed on ice, 100μL chloroform was added, vortexed for 30 seconds, and centrifuged at 12,000*g* for 15 minutes at 4°C. After centrifugation the aqueous phase was transferred to a QIAGEN gDNA eliminator column and centrifuged at 10,000*g* for 1 minute at room temperature. The flow-through was mixed with an equal volume of 70% EtOH and transferred to an RNEasy mini spin column. The manufacturer protocol for the RNEasy Plus Mini Kit (QIAGEN) was followed for the rest of the procedure. cDNA was synthesized using the high Capacity cDNA reverse transcription (Applied Biosystems). qRT-PCR was performed using the SYBR Green intercalating dye (ThermoFisher Scientific). Expression was normalized to 18S ribosomal RNA and relative gene expression was calculated using $2^{-\Delta\Delta CT}$ method.

**Immunofluorescence Staining**

*Staining on glass coverslips*

Cells were grown on Matrigel-coated 8mm or 12mm diameter circular glass coverslips. Cultures were fixed for 20 minutes in 4% paraformaldehyde (Electron Microscopy Sciences) at room temperature. Coverslips were then washed three times with PBS followed by 15 minutes permeabilization with 0.2% Triton X-100 in PBS. Subsequently the coverslips were washed three times in washing buffer (0.05% Tween-20 in PBS) for 5 minutes and blocked for 20 minutes in 200 μl wash buffer plus 5% normal donkey serum (Jackson ImmunoResearch Laboratories). The primary antibodies were diluted in 5% normal donkey serum in PBS and incubated with the tissues 1 hour at room temperature followed by three washes in wash buffer for 5 minutes each. The secondary antibodies were diluted in 5% normal donkey serum in PBS and incubated with the tissues 1 hour at room temperature followed by three washes in wash buffer for 5 minutes each. Afterwards, the coverslips were mounted on microscopy glass slides using ProLong Diamond Antifade (Life Technologies), cured overnight at room temperature and then sealed with nail polish. Antibody list is in the STAR methods.

*Paraffin Embedded Sectioning and Staining*

Tissue samples harvested from DesLO implanted mice washed in PBS and incubated in 4% paraformaldehyde at 4°C overnight. The samples were washed 3 times with PBS, transferred to a 70% ethanol solution, and delivered to Mayo Clinic Histology Core Laboratory for paraffin embedding and sectioning (4-5um slices). The returned slides were submerged 3 times with fresh xylene for 5 minutes then washed twice for 10 minutes in 100% ethanol, followed by twice for 10 minutes in 95% ethanol, then twice more in deionized (DI) water. Antigen retrieval was performed by submerging the slides in 1x citrate buffer (Abcam) and kept at the brink of boiling for 15 minutes in a microwave oven. The slides were then washed 3 times in DI water for 5 minutes, then once in PBS. Samples were blocked and permeablized in PBS + 8% donkey serum + 0.2% TritonX-100 for 2 hours at room temperature. Primary antibodies were incubated with the samples overnight at 4°C, rinsed 3 times in DI water, then washed 15 minutes in PBS. The secondary antibodies were added in PBS + 2% donkey serum for 2 hours at room temperature. Tissues were then counterstained with either DAPI or Hoechst, rinsed 3 times in DI water, washed for 15 minutes in PBS, and sealed with a glass coverslip for imaging using either Diamond Antifade (Thermo Scientific) or Vectorshield (Vector Labs).

**Indocyanine Green, Oil Red O, and Periodic acid-Schiff Staining**

Indocyanine green (Cat. #21980-100MG-F) was added to cultures at 1mg/mL for 30 minutes at 37°C, washed 3x in culture medium, and imaged. For Oil Red O (Cat. #O1391-250ML), cultures were fixed in 4% paraformaldehyde for 20 minutes at room temperature, rinsed 3x with DI water, treated with 60% isopropanol for 5 minutes, incubated with Oil Red O for 15 minutes, rinsed with DI water until there was no detectable red solution in the wash water, and then imaged. For Periodic acid-Schiff (Cat. #395B-1KT) cultures were fixed in 4% paraformaldehyde for 20 minutes, rinsed 3x with 1X PBS, incubated in periodic acid for 5 minutes, washed 3x with DI water, incubated for 15 minutes Schiff's reagent, rinsed 3x with DI water, and imaged.

**Image Acquisition and Processing**

Images were acquired using the Leica DMi8 automated scanning microscope or Leica TCS SP5 confocal microscope, and processed using ImageJ software (NIH). AngioTool software (NIH) was used to quantify the vascular formation metrics from single channel grayscale images of vascular stains (CD31 staining). 3D reconstructions were generated using the Leica TCS SP5 confocal microscope to generate z-stacks spanning ~200um deep into the tissues, and using Image J to construct a 3D volume from the stacks.

**HNF4A$^+$ Quantification**

DesLO were stained for HNF4A, imaged using a DMi8 scanning fluorescent microscope, and quantified using ImageJ with the following pipeline: Threshold > Make Binary > Fill Holes > Watershed > Analyze Particles. Averaged results were used for normalization of DesLO in ELISA assays, CYP3A4 activity and urea when expressed in "…/M-cells".

**Enzyme-linked Immunosorbent Assays (ELISA)**

Samples were assayed for AAT (Genway Biotech), albumin (Bethyl Labs), C3 (Immunology Consultants Laboratory), and ANGPTL3 (RayBiotech) using commercially available ELISA kits. Cultures being assayed for human albumin were switched to daily feeding with William's medium E prior to sampling for measurement with ELISA due to the presence of human albumin in APEL medium. Sample dilutions were optimized to attain detection in the linear range of the standard curves for each individual assay.

**Cytochrome P450 Activity Assays**

The CYP3A4 and CYP2C19 activity assays were performed according to the manufacturer's instructions for Cell-based assays (P450-Glo Assays, Promega) and analyzed with a luminometer. CYP3A4 activity was calculated using a beetle luciferin standard curve measured with the samples.

**Total Bile Acid and Urea Assays**

The total bile acid assay (Cell Biolabs) and Urea assay (Cell Biolabs) were used to measure the bile acid and urea concentrations in cultures. The cell supernatant was assessed at multiple dilutions to optimize detection, and the assays were performed according to the manufacturer's protocol.

**Mouse Studies and Implants**

For implantation, engineered hiPSCs were seeded on 50% Matrigel 3-D gels (1:1 RGF Matrigel and DMEM/F12) in 48-well plates to allow contraction of cell layer into a dense tissue or adhered to Matrigel-coated Cytodex3 microcarrier beads (Sigma, 1000:1 cell to bead ratio) in 96-well round bottom plates instead of Matrigel monolayer coated tissue culture polystyrene. DesLO tissues were implanted either beneath the renal capsule or on the mesentery, and secured in place using a fibrin gel supplemented with 50 ng/mL hHGF and 20 ng/mL hVEGF-165 to promote survival and angiogenesis of the tissue for integration with host vasculature. For survival studies, FRGN (Yecuris) mice between 8 and 12 weeks old were used. For the TK-NOG studies male mice between 8 and 12 week old were used, and ganciclovir (50 mg/kg, IP), which is innocuous to normal human and mouse cells, was administered to induce damage of the mouse liver parenchymal cells at day 7, 10, and 20 after implantation of 10-12 DesLOs (~3 million cells) onto the mesentery. For FRGN studies, nitisinone (NTBC) was removed from the drinking water directly after implantation of up to 5 DesLOs under the renal capsule, or 10-12 DesLOs onto the mesentery. The NTBC treatment was cycled once, being added to the drinking water at 16mg/L after 14 days for 5 days, and then removed permanently. For control experiments, sham surgeries were performed identical to the implants but with no implanted tissues. Primary human hepatocyte (3 lots) and iHEP (3 lots) controls were generated by embedding 3 million cells in Matrigel, securing them to the mesentery with the same fibrin gel and growth factor recipe as DesLO, and maintaining under the same conditions as the DesLO-implanted mice. Blood was drawn from mice using cheek puncture. Kaplan-Meier survival analysis was performed with Prism 8 (GraphPad Software Inc.).

**Bulk RNA sequencing of FeLO and DesLO tissues**

RNA was extracted from samples as described above and sent to the UCLA Technology Center for Genomics and Bioinformatics for library preparation and sequencing. Libraries for RNA-Seq were prepared with KAPA Hyper Stranded RNA-Seq Kit. The workflow consists of mRNA enrichment, cDNA generation, and end repair to generate blunt ends, A-tailing, adaptor ligation and PCR amplification. Different adaptors were used for multiplexing samples in one lane. Sequencing was performed on Illumina NextSeq500 for a single read 75bp run. Data quality check was done on Illumina SAV. Demultiplexing was performed with Illumina Bcl2fastq2 v 2.17 program. A raw FASTQ quality check was performed using FASTQC (http://www.bioinformatics.babraham.ac.uk/projects/fastqc). Reads were then mapped to the latest UCSC transcript set using Bowtie2 version 2.1.0 (Langmead and Salzberg, 2012) and the gene expression level was estimated using RSEM v1.2.15 (Li and Dewey, 2011). EdgeR's (Robinson et al., 2010) TMM (trimmed mean of M-values) algorithm was used to normalize the gene expression data. Reads were then mapped to the latest UCSC genome set using Bowtie2 and Tophat (Trapnell et al., 2009). The resulting BAM file allowed for collection of information on the alignment via PicardTools (http://broadinstitute.github.io/picard/) CollectRNASeqMetrics program. A Genebody analysis was performed using the ngsplot (Shen et al., 2014) toolkit.

**Enrichr Analysis and Heatmaps for Pathways**

Differential expression data was generated from the TMM normalized reads (counts per million, CPM) by dividing the average counts of each sample over the average counts for the reference condition for each gene. The resulting fold change list was used to generate a list of genes for each condition with at least two reads and ≥ 2 fold change over the reference condition. The genes from each resulting list were imported into the EnrichR web browser application, and the results used to generate alignment scoring and significance data for cell, pathway, function, and ontology analyses. For scRNA-Seq data gene lists were generated using Seurat for each cluster including genes with adj. p-values < .05 and fold change expression of at least 1.6-fold over the average gene expression of the other clusters. The lists were then submitted to EnrichR for enrichment analysis and results displayed using Graphpad Prism 8.

Genes from liver pathways illustrated using heatmaps were defined using available pathway lists from Enrichr (Chen et al., 2013; Kuleshov et al., 2016) and from additional publications. Genes were binned according to enrichment at each stage of DesLO engineering (F = untransduced FeLO, P = PROX1, PA = PROX1 + ATF5, D (DesLO) = PROX1 + ATF5 + SynTF(*CYP3A4*)) by fold change relative to untransduced FeLO for genes with at least two reads. Bins are defined as genes that have over two-fold enrichment for P, genes that have over two-fold enrichment for PA and not P, and genes that have over two-fold enrichment for D and not P or PA. Up to 15 genes for P enrichment, 5 genes for PA enrichment, and 5 genes for D enrichment were shown per pathway in heatmaps (depicted as $log_2$ fold change expression over FeLO), with occasional addition of select genes of interest from the pathway in order to show the expression of key genes of interest to the community. Due to significant overlap of gene lists included, the Fat metabolism and Cholesterol and lipid metabolism pathway results were combined after binning and displayed together as "Fat and lipid metabolism". The same was done for the combination of the Bile Acid Secretion and FXR Signaling pathways. In the case of no expression in FeLO for genes of interest, FeLO expression was assigned the lowest detectable expression value for comparison.

**Assay for Transposase-Accessible Chromatin Using Sequencing (ATAC-Seq)**

Day 17 DesLO were in acquired in single cell suspension by incubation with trypsin for 10 minutes at 37°C, followed by gentle pipetting using a serological pipette to dislodge and dissociate aggregates. Cells were frozen in mFresR cryopreservation medium. ATAC-Seq library preparation and sequencing reactions were conducted at GENEWIZ, LLC. (South Plainfield, NJ, USA). Live cell samples were thawed, washed, and treated with DNAse I (Life Tech, Cat. #EN0521) to remove genomic DNA contamination. Live cell samples were quantified and assessed for viability using a Countess Automated Cell Counter (ThermoFisher Scientific, Waltham, MA, USA). After cell lysis and cytosol removal, nuclei were treated with Tn5 enzyme (Illumina, Cat. #20034197) for 30 minutes at 37°C and purified with Minelute PCR Purification Kit (QIAGEN, Cat. #28004) to produce tagmented DNA samples. Tagmented DNA was barcoded with Nextera Index Kit v2 (Illumina, Cat. #FC-131-2001) and amplified via PCR prior to a SPRI Bead cleanup to yield purified DNA libraries. The sequencing libraries were clustered on a single lane of a flow cell. After clustering, the flow cell was loaded on the Illumina HiSeq instrument (4000 or equivalent) according to manufacturer's instructions. The samples were sequenced using a 2x150bp Paired End (PE) configuration. Image analysis and base calling were conducted by the HiSeq Control Software (HCS). Raw sequence data (.bcl files) generated from Illumina HiSeq was converted into FASTQ files and de-multiplexed using Illumina's bcl2fastq 2.17 software. One mismatch was allowed for index sequence identification.

*ATAC-seq Data Pre-processing:*

ATAC-seq data downstream of FASTQ file generation was analyzed by the PLRC Genomics and Systems Biology Core. Raw ATAC-seq reads first went through the pipeline of FastQC (http://www.bioinformatics.babraham.ac.uk/projects/fastqc) for quality control. Reads with low sequencing quality and adapter sequences were trimmed out by software Trimmomatic (Bolger et al., 2014). Then the surviving reads were aligned to reference genome hg19 by Burrows-Wheeler Aligner (Li and Durbin, 2009). Reads with low mapping quality were filtered out and only primary alignments were kept by SAMtools (Li et al., 2009). Duplicate reads were marked by Picard tool http://broadinstitute.github.io/picard/ . Once the alignment was done, MACS (Zhang et al., 2008) was applied for peak calling.

*Differential Binding Site Analysis*

In order to compare the differential binding sites between pairwise conditions across hiPSC, FeLO and DesLO, peaks called from each individual sample were pooled together will common peak regions. Read count for each pooled peak region within each sample were quantified by R package 'DiffBind' http://bioconductor.org/packages/release/bioc/html/DiffBind.html. Based on peak enrichment, correlation plot and PCA plot were generated to visualize the correlation/ similarity across the samples. Differential binding sites test were performed by R package 'DESeq2' (Love et al., 2014). By cutoff of FDR=5% and absolute $\log_2$(fold change) greater than 1, top peaks were selected, representing genomic regions that are more open in one condition compared with the other one. Selected peak regions were visualized by tool IGV (Robinson et al., 2011).

#### Single Nucleotide Polymorphism (SNP) Analysis

Genomic data for the hiPSC line used in this study was obtained from the personal genome project (https://my.pgp-hms.org). SNPs were extracted for *CYP3A4* region (chr7:99354583-99381811) by reference genome hg19.

#### 10x Genomics Sample Preparation for Next-generation Sequencing

Samples were prepared as described by the 10x Genomics Single Cell 3′ v2 Reagent Kit user guide. Day 17 DesLO were acquired in single cell suspension by incubation with trypsin for 10 minutes at 37°C , followed by gentle pipetting using a serological pipette to dislodge and dissociate aggregates. Two washes in PBS -/- + 0.04% BSA were performed with the cells re-suspended at a final concentration of 1000 cells/µL in PBS -/- + 0.04% BSA. A live cell count was performed with a hemocytometer using Trypan Blue to identify dead cells. Following counting, cells and 10x Genomics reagents were loaded into the single cell cassette, with a target of 6000 single cells for analysis, accounting for predicted cell loss and doublets as laid out in the user guide for the Chromium Single-Cell 3' Reagent Kit (10x Genomics). After generation of GEMs, the cDNA library was prepared by ASU Genomics Core staff following the appropriate steps determined by the 10x Genomics user guide. Libraries were sent to Novogene for sequencing by an Illumina HiSeq X for an intended read depth of 100,000 reads per cell with 150 bp paired end reads. Our downstream analysis from the sequencing data estimates the actual number of cells sequenced was between 5000 and 7000 which would yield >80k reads per cell. The 10x Genomics CellRanger pipeline was used to align reads to the reference genome (GRCh38.84) appended with transgene sequences, to assign reads to individual cells, and to estimate gene expression based on UMI counts (Zheng et al., 2017).

Single cell data was excluded based on high mitochondrial genome transcript ratio and either high or low feature or UMI counts. Genes with UMI counts in fewer than 5 cells were removed from consideration. For scRNA-Seq data processing and cluster analysis using Seurat (Butler et al., 2018; Satija et al., 2015), we used the following general standardized pipeline for processing of the Cell Ranger output: Normalization, feature selection, scaling (including cell cycle regression), principal component analysis (PCA), and clustering. Elbow plots and permuted p-values were used to assist in determining the optimal number of principal components (PCs) needed to summarize the datasets without losing a significant amount of variation. The quality of a range of clustering resolution values were assessed using enrichment of cluster marker genes (genes differentially upregulated in a given cluster relative to all other clusters) with liver cell type-specific genes. As a quality check, PC and resolution metrics were modulated to yield fewer or additional clusters to confirm that chosen parameters resulted in the most biologically relevant clustering. Visualization was achieved by the use of tSNE plots identifying cells, clusters, and selected gene expression in each cell, as well as heatmaps and violin plots showing the expression level of genes by cluster. Expression per cell by cluster was also extracted and visualized using Graphpad Prism 8.

#### Comparison of Endothelial- and Stellate-like Cell Clusters with Single Cell Analysis of Primary Liver Samples

The top 25 differentially expressed genes in LSEC zone1, LSEC zone 2, periportal endothelial, and stellate cell clusters from the available primary liver scRNA-Seq data of MacParland et al., 2018, were cross referenced against ≥1.5-fold upregulated (compared to average expression of the genes across all clusters) genes from the ELC and SLC clusters from the DesLO scRNA-Seq. The top 25 genes from MacParland et al. were displayed in a binary heatmap for LSEC zone 1, LSEC zone 2, periportal endothelial, and stellate cell clusters along with the DesLO ELC and SLC clusters.

**CellNet Analysis of FeLO Time Lapse and DesLO Bulk RNA**

We used the bulk RNA-Seq CellNet (Radley et al., 2017) pipeline to quantify gene expression estimates as previously described, aligning to reference genome GRCh38. Classification and network analysis were performed using the human cnProc cnProc_HS_RS_Apr_05_2017, which is trained on 15 cell and tissue types from mostly adult sources[11]. Source code are available through GitHub (https://github.com/pcahan1/CellNet).

For meta-analysis, we used CellNet to quantify gene expression, and performed classification and gene regulatory network analysis for bulk RNA sequencing samples from past studies (Akbari et al., 2019; Asai et al., 2017; Du et al., 2014; Ouchi et al., 2019) in the same manner as we did with our FeLO and DesLO bulk RNA seq samples. We compared the classification scores and liver gene regulatory network status across all studies using classification heatmaps and bar plots.

**KeyGenes Analysis**

Raw gene pseudocounts produced by SALMON (Patro et al., 2017) were run through the KeyGenes algorithm (https://github.com/DavyCats/KeyGenes). Human fetal (excluding the maternal endometrium) (Roost et al., 2015) and adult data (Fagerberg et al., 2014); Illumina Body Map 2.0) were used as training data.

**QUANTIFICATION AND STATISTICAL ANALYSIS**

For studies in which statistical analyses were performed, at least three biologically independent replicates were used unless otherwise indicated. Statistical comparisons of three or more conditions were performed using one-way ANOVA and multiple comparisons testing on means using Tukey's method with a $p$-value $< 0.05$ as the threshold for significance. This method was used where significance has been reported without indicating method. Otherwise, for comparisons between two conditions, multiple groups, or survival curves, one or two tailed t-tests, two-way ANOVA with multiple comparison test, or Mantel-Cox tests were performed with a $p$-value $< 0.05$ as the threshold for significance and are indicated as such. Logrank p-values for the FRGN survival study were determined using the Mantel-Cox test.

### KEY RESOURCES TABLE

| REAGENT or RESOURCE | SOURCE | IDENTIFIER |
|---|---|---|
| Antibodies | | |
| AAT | R&D | Cat# AF1268; RRID:AB_354707 |
| Albumin | Bethyl | Cat# A80-229A, RRID:AB_67018 |
| CD146 | R&D | Cat# AF932, RRID:AB_355721 |
| CD31 | Cell Signaling Technologies | Cat# 3528, RRID:AB_2160882 |
| CD31 | Cell Signaling Technologies | Cat# 77699, RRID:AB_2722705 |
| CD34 | Abcam | Cat# ab81289, RRID:AB_1640331 |
| CDX2 | BioGenex | Cat# AM392, RRID:AB_2650531 |
| CEBPA | R&D | Cat# AF7094, RRID:AB_10973004 |
| CK18 | Santa Cruz Biotechonology | Cat# sc-32329, RRID:AB_627849 |
| Desmin | Santa Cruz Biotechonology | Cat# sc-7559, RRID:AB_639081 |
| Desmin | Santa Cruz Biotechonology | Cat# sc-23879, RRID:AB_627416 |
| E-Cadherin | Cell Signaling Technologies | Cat# 3195, RRID:AB_2291471 |
| EPCAM | Cell Signaling Technologies | Cat# 36746, RRID:AB_2799105 |
| ERG | Abcam | Cat# ab92513, RRID:AB_2630401 |
| FOXA2 | R&D | Cat# AF2400, RRID:AB_2294104 |
| GFP | Abcam | Cat# ab13970; RRID:AB_300798 |
| HNF4A | Cell Signaling Technologies | Cat# 3113; RRID:AB_2295208 |
| NANOG | R&D | Cat# AF1997; RRID:AB_355097 |
| NANOG | Cell Signaling Technologies | Cat# 3580; RRID:AB_2150399 |
| Nestin | Santa Cruz Biotechonology | Cat# sc-23927; RRID:AB_627994 |
| SOX17 | R&D | Cat# AF1924; RRID:AB_355060 |
| T/Brachyury | R&D | Cat# AF2085; RRID:AB_2200235 |
| AF-488 anti-chicken | Jackson Immunoresearch Laboratories | Cat# 703-545-155; RRID:AB_2340375 |
| AF-594 anti-chicken | Jackson Immunoresearch Laboratories | Cat# 703-585-155; RRID:AB_2340377 |
| AF-488 anti-goat | Thermo Fisher | Cat# A-11055; RRID:AB_2534102 |
| AF-594 anti-goat | Thermo Fisher | Cat# A-11058; RRID:AB_2534105 |
| AF-647 anti-goat | Thermo Fisher | Cat# A-21447; RRID:AB_141844 |
| AF-488 anti-rabbit | Thermo Fisher | Cat# A-21206; RRID:AB_2535792 |
| AF-594 anti-rabbit | Jackson Immunoresearch Laboratories | Cat# 711-585-152; RRID:AB_2340621 |
| AF-647 anti-rabbit | Thermo Fisher | Cat# A-31573; RRID:AB_2536183 |
| AF-488 anti-mouse | Thermo Fisher | Cat# A-21202; RRID:AB_141607 |
| AF-594 anti-mouse | Jackson Immunoresearch Laboratories | Cat# 715-585-151; RRID:AB_2340855 |
| AF-647 anti-mouse | Thermo Fisher | Cat# A-31571; RRID:AB_162542 |
| AF-594 anti-Sheep | Thermo Fisher | Cat# A-11016; RRID:AB_2534083 |
| AF-647 anti-Sheep | Thermo Fisher | Cat# A21448; RRID:AB_1500712 |
| APC-CD34 | Miltenyi | Cat# 130-090-954, RRID:AB_244349 |
| PE-CD146 | Miltenyi | Cat# 130-097-939, RRID:AB_2660768 |
| Bacterial and Virus Strains RL | | |

| | | |
|---|---|---|
| NEB 5 alpha competent E. coli | New England Biolabs | Cat# C2987H |
| Biological Samples RL | | |
| Human Adult Liver Total RNA | Cell Applications, Inc | Cat# 1H21-50 |
| Human Liver Total RNA | Takara Bio | Cat# 636531 |
| Human Adult Liver Total RNA | Thermo Fisher Scientific | Cat# AM7960 |
| Chemicals; Peptides; and Recombinant Proteins | | |
| Axitinib | Cell Signaling Technology | Cat# 12961S |
| Phosphate buffered saline | Corning | Cal# 21-040-CV |
| Polybrene | Millipore-Sigma | Cat# TR-1003-G |
| Thawing/Plating Cocktail A | Thermo Fisher Scientific | Cat# CM3000 |
| Cell Maintenance Cocktail B | Thermo Fisher Scientific | Cat# CM4000 |
| Rat tail collagen 1 | Thermo Fisher Scientific | Cat# A1048301 |
| GW4064 | Sigma Aldrich | Cat# G5172-5MG |
| Chenodeoxycholic acid | Cayman Chemical | Cat# 10011286 |
| Human recombinant FGF19 | Peprotech | Cat# 100-32 |
| human recombinant TGFβ1 | Peprotech | Cat# 100-21 |
| Human recombinant HGF | Peprotech | Cat# 100-39H |
| Human recombinant VEGF-165 | Peprotech | Cat# 100-20 |
| Dimethylsulfoxide | Sigma Aldrich | Cal# D2650-100ML |
| TRIzol Reagent | Thermo Fisher Scientific | Cat# 15596018 |
| Y-27632 Dihydrochloride | Stem Cell Technologies | Cat# 72305 |
| Puromycin Dihydrochloride | Sigma Aldrich | Cat# P8833 |
| Doxycycline hyclate | Sigma Aldrich | Cat# D9891 |
| Chloroform | Sigma Aldrich | Cat# 288306 |
| DMEM | Gibco | Cat# 11960069 |
| DMEM/F12 | Gibco | Cat# 11320082 |
| Williams E Medium | Thermo Fisher Scientific | Cat# A1217601 |
| mTeSR 1 | Stem Cell Technologies | Cat# 85850 |
| mFresR | Stem Cell Technologies | Cat# 05855 |
| STEMdiff APEL | Stem Cell Technologies | Cat# 28995 |
| Accutase | Stem Cell Technologies | Cat# 07922 |
| hESC qualified matrigel | Corning | Cat# 354277 |
| GFR, LDEV free matrigel | Corning | Cat# 354230 |
| Lipofectamine 3000 Transfection Reagent | Thermo Fisher Scientific | Cat# L3000001 |
| Super PiggyBac Transposase Expression Vector | System Biosciences | Cat# PB210PA-1 |
| RNeasy Plus Mini Kit | QIAGEN | Cat# 74134 |
| AllPrep DNA/RNA Mini Kit | QIAGEN | Cat# 80204 |
| SYBR green power up | Thermo Fisher Scientific | Cat# A25742 |
| Normal donkey serum | Jackson Immunoresearch Laboratories | Cat# 017-000-001 |
| Prolong Diamond Antifade | Thermo Fisher Scientific | Cat# P36970 |

| | | |
|---|---|---|
| Xylenes | Fisher Chemical | Cat# X3S-4 |
| Anhydrous Ethanol | Fisher Chemical | Cat# A405P-4 |
| DAPI | Thermo Fisher Scientific | Cat# 62248 |
| Heocsht 33342 | Thermo Fisher Scientific | Cat# H3570 |
| Indocyanine Green | Sigma | Cat# 21980-100MG-F |
| Oil Red O | Sigma | Cat# O1391-250ML |
| Cytodex3 Microcarrier Beads | Sigma | Cat# C3275-10G |
| Anti-TRA-1-60 MicroBeads, human | Miltenyi Biotec | Cat# 130-100-832 |
| Anti-CD34 MicroBeads, human | Miltenyi Biotec | Cat# 130-046-702 |
| Anti-CD146 MicroBeads, human | Miltenyi Biotec | Cat# 130-093-596 |
| Nitisinone (NTBC) | Sigma | Cat# PHR1731-1G |
| Ganciclovir Sodium Salt | Santa Cruz Biotechnology | Cat# sc-394139B |
| Critical Commercial Assays | | |
| Bethyl Albumin ELISA KIT | Bethyl | Cat# E80-129 |
| SERPINA1 (AAT) ELISA | Genway Biotech | Cat# GWB-1F2730 |
| Fibrinogen ELISA | Genway Biotech | Cat# GWB-C5E724 |
| C3 ELISA | Immunology Consultants Laboratory | Cat# E-80C3 |
| ANGPTL3 ELISA | RayBiotech | Cat# ELH-ANGPTL3-1 |
| Periodic Acid-Schiff (PAS) Kit | Millipore-Sigma | Cat# 395B-1KT |
| Total bile acid | Cell Biolabs | Cat# STA-631 |
| Urea assay | Bioassay Systems | Cat# DIUR-100 |
| qPCR Lentivirus titration kit | Applied Biological Materials | Cat# LV900 |
| CYP2C19 P450-Glo™ assay | Promega | Cat# V8881 |
| CYP3A4 P450-Glo™ Assay | Promega | Cat# V9001 |
| Deposited Data | | |
| Bulk RNA sequencing of FeLO, engineered FeLO conditions and DesLO | This paper | |
| Day 2, 6 liver bud, cryopreserved human hepatocytes, and whole liver RNA sequencing data | Asai et al., 2017 | GEO Accession GSE85223: GSM2262397, GSM2262398, GSM2262403, GSM2262404, GSM2262407, and GSM2262406 |
| FeLO and DesLO day 17 single cell RNA sequencing | This paper | |
| Human liver organoids in differentiation medium RNA sequencing data | Akbari et al., 2019 | GEO Accession GSE130075: GSM3731529 and GSM3731530 |
| Human liver organoids D25 | Ouchi et al., 2019 | GEO Accession GSE130075: GSM3731529 and GSM3731530 |
| Fibroblast transdifferentiated hepatocytes and freshly isolated hepatocytes | Du et al., 2014 | GEO Accession GSE54066: GSM1306654 and GSM1306653 |
| Adult human liver single cell RNA sequencing | MacParland et al., 2018 | GEO Accession GSE115469 |
| Experimental Models: Cell Lines | | |

| | | |
|---|---|---|
| HEK293FT | Thermo Fisher Scientific | Catalog# R70007 |
| PGP1 rttA3 TRE GATA6-2A-EGFP | This paper | |
| Primary Human Hepatocytes | Thermo Fisher Scientific | Catalog# HMCPMS; lot# HU8295 |
| Primary Human Hepatocytes | MGH Cell Resource Core | lot# HH-083 |
| Primary Human Hepatocytes | Lonza | Catalog# HMCPM; lot# HUM4012 |
| Primary Human Hepatocytes | Lonza | Catalog# HUCPG; lot# HUM17299A, lot# HUM180851 |
| Primary Human Hepatocytes | Zenbio | Catalog# HUCPG; lot# ZBH1989-P |
| Cellartis Enhanced hiPS-HEP v2 | Takara Bio Inc | Catalog# Y10133, Y10134 |
| iCell Hepatocytes | Stem Cell Technologies | Catalog# R1104 |
| Experimental Models: Organisms/Strains | | |
| Mouse: TK-NOG, NOD.Cg-Prkdcscid Il2rgtm1Sug Tg(Alb-TK)7-2/ShiJic | Taconic Biosciences | Cat# 12907-M |
| Mouse: FRG® KO on NOD | Yecuris Corporation | Cat# 10-0008 |
| Oligonucleotides | | |
| CYP3A4 sgRNA Target Sequence 1: ACTCAAAGGAGGTCAGTGAG | This paper | N/A |
| CYP3A4 sgRNA Target Sequence 2: TGATTCTTTGCCAACTTCCA | This paper | N/A |
| Primers for qPCR, See Table 1 | This paper | N/A |
| Primer: NANOG Forward: ACAACTGGCCGAAGAATAGCA | This paper | N/A |
| Primer: NANOG Reverse: GGTTCCCAGTCGGGTTCAC | This paper | N/A |
| Primer: SOX2 Forward: GCTTAGCCTCGTCGATGAAC | This paper | N/A |
| Primer: SOX2 Reverse: AACCCCAAGATGCACAACTC | This paper | N/A |
| Primer: TBXT Forward: ACAAAGAGATGATGGAGGAACCCG | This paper | N/A |
| Primer: TBXT Reverse: AGGATGAGGATTTGCAGGTGGACA | This paper | N/A |
| Primer: FOXA2 Forward: CGACTGGAGCAGCTACTATGC | This paper | N/A |
| Primer: FOXA2 Reverse: TACGTGTTCATGCCGTTCAT | This paper | N/A |

| | | |
|---|---|---|
| Primer: CD34 Forward: ACCAGAGCTATTCCCAAAGACC | This paper | N/A |
| Primer: CD34 Reverse: TGCGGCGATTCATCAGGAAAT | This paper | N/A |
| Primer: DES Forward: CTGAGCAAAGGGGTTCTGAG | This paper | N/A |
| Primer: DES Reverse: ACTTCATGCTGCTGCTGTGT | This paper | N/A |
| Primer: HNF4a Forward: CGTCATCGTTGCCAACACAAT | This paper | N/A |
| Primer: HNF4a Reverse: GGGCCACTCACACATCTGTC | This paper | N/A |
| Primer: ALB Forward: GCCTTTGCTCAGTATCTT | This paper | N/A |
| Primer: ALB Reverse: AGGTTTGGGTTGTCATCT | This paper | N/A |
| Primer: ASGR1 Forward: ATGACCAAGGAGTATCAAGACCT | This paper | N/A |
| Primer: ASGR1 Reverse: TGAAGTTGCTGAACGTCTCTCT | This paper | N/A |
| Primer: NR0B2 Forward: GTGCCCAGCATACTCAAGAAG | This paper | N/A |
| Primer: NR0B2 Reverse: TGGGGTCTGTCTGGCAGTT | This paper | N/A |
| Primer: CYP7A1 Forward: GAGAAGGCAAACGGGTGAAC | This paper | N/A |
| Primer: CYP7A1 Reverse: GGATTGGCACCAAATTGCAGA | This paper | N/A |
| Primer: TIMP1 Forward: CTTCTGCAATTCCGACCTCGT | This paper | N/A |
| Primer: TIMP1 Reverse: ACGCTGGTATAAGGTGGTCTG | This paper | N/A |
| Primer: LOX Forward: CGCTGTGACATTCGCTACACAGGAC | This paper | N/A |
| Primer: LOX Reverse: CATTGGGAGTTTTGCTTTGCCTTCT | This paper | N/A |
| Primer: COL1A1 Forward: TCCCACCAATCACCTGCGTACA | This paper | N/A |

| | | |
|---|---|---|
| Primer: COL1A1 Reverse: CGCCGGTGGTTTCTTGGTCG | This paper | N/A |
| Primer: FGF19 Forward: CGGAGGAAGACTGTGCTTTCG | This paper | N/A |
| Primer: FGF19 Reverse: CTCGGATCGGTACACATTGTAG | This paper | N/A |
| Primer: MLXIPL Forward: AACCGGCGTATCACACACAT | This paper | N/A |
| Primer: MLXIPL Reverse: TGTCTTCTGCAGCGTGGTAG | This paper | N/A |
| Primer: MCAM Forward: GGTCGCTACCTGTGTAGGGA | This paper | N/A |
| Primer: MCAM Reverse: TGGACCCGGTTCTTCTCCT | This paper | N/A |
| Primer: PECAM1 Forward: TGTATTTCAAGACCTCTGTGCACTT | This paper | N/A |
| Primer: PECAM1 Reverse: TTAGCCTGAGGAATTGCTGTGTT | This paper | N/A |
| Primer: ACTA2 Forward: CGGCTTTGCTGGGGACGAT | This paper | N/A |
| Primer: ACTA2 Reverse: CAGGGGCAACACGAAGCTCAT | This paper | N/A |
| Primer: NES Forward: GAAGGGCAATCACAACAGGTG | This paper | N/A |
| Primer: NES Reverse: GGGGCCACATCATCTTCCA | This paper | N/A |
| Primer: RNA18S5 | Integrated DNA Technologies | Hs.PT.39a.22214856.g |
| Primer: NR1H4 | Integrated DNA Technologies | Hs.PT.58.18707722 |
| Primer: ATF5 | Integrated DNA Technologies | Hs.PT.58.25040399 |
| Primer: PROX1 | Integrated DNA Technologies | Hs.PT.58.2987292 |
| Primer: CYP3A4 | Integrated DNA Technologies | Hs.PT.58.1272782 |
| Primer: GATA6 | Integrated DNA Technologies | Hs.PT.58.38396504 |
| Primer: G6PC | Integrated DNA Technologies | Hs.PT.58.46489349 |
| Primer: CREB3L3 | Integrated DNA Technologies | Hs.PT.58.27018013 |
| Recombinant DNA | | |
| psPax2 | Trono Lab Packaging and Envelope Plasmids (Unpublished) | Addgene Plasmid# 12260 |
| pCMV-VSV-G | Stewart et al 2003 | Addgene Plasmid# 8454 |
| MS2-P65-HSF1-GFP | Konermann et al 2014 | Addgene Plasmid# 61423 |
| U6-sgRNA-MS2 | Konermann et al 2014 | Addgene Plasmid# 61424 |

| | | |
|---|---|---|
| Cas9M4-VP64 | Mali et al 2013 | Addgene Plasmid# 47319 |
| pENTR_L1_hGATA6-2A-EGFP_L2 | Guye et al 2016 | N/A |
| PB-TAG-ERP2 | Kim et al 2016 | Addgene Plasmid#80479 |
| PROX1 transcript variant 2 cDNA | Origene | Cat# RC200081 |
| ATF5 transcript variant 1 cDNA | Genecopoeia | Cat# F0925 |
| CREB3L3 complete CDS cDNA | DNASU Plasmid Repository | (Clone ID# HsCD00080068) |
| MLXIPL transcript variant 1 cDNA | DNASU Plasmid Repository | Clone ID# HsCD00820703 |
| Promoter: AAT Sequence: AGGTATCTTGCTACCAGTG GAACAGCCACTAAGGATTCT GCAGTGAGAGCAGAGGGCC AGCTAAGTGGTACTCTCCCA GAGACTGTCTGACTCACGC CACCCCCTCCACCTTGGAC ACAGGACGCTGTGGTTTCT GAGCCAGGTACAATGACTC CTTTCGGTAAGTGCAGTGG AAGCTGTACACTGCCCAGG CAAAGCGTCCGGGCAGCGT AGGCGGGCGACTCAGATCC CAGCCAGTGGACTTAGCCC CTGTTTGCTCCTCCGATAAC TGGGGTGACCTTGGTTAATA TTCACCAGCAGCCTCCCCC GTTGCCCCTCTGGATCCAC TGCTTAAATACGGACGAGG ACAGGGCCCTGTCTCCTCA GCTTCAGGCACCACCACTG ACCTGGGACAGTGAATCGT AAGTGCTT | This paper | N/A |
| Software and Algorithms | | |
| CellNet | P. Cahan Lab | https://github.com/pcahan1/CellNet |
| R version 3.6.2 | The Comprehensive R Archive Network | https://cran.r-project.org/ |
| R Studio version 1.2.5033 | RStudio Inc. | https://rstudio.com/ |
| Cell Ranger | 10x Genomics | https://www.10xgenomics.com/ |
| Graphpad Prism 8 | Graphpad Software Inc. | https://www.graphpad.com/ |
| FlowJo | Becton, Dickson and Co. | https://www.flowjo.com/ |
| IGV (version 2.7.2) | Broad Institute | https://software.broadinstitute.org/software/igv/ |
| Angiotool (version 0.6a) | NIH National Cancer Institute Center for Cancer Research | https://ccrod.cancer.gov/confluence/display/ROB2/Downloads |
| ImageJ (version 1.48v) | NIH | https://imagej.nih.gov/ij/ |
| Enrichr | A. Ma'ayan Lab | https://amp.pharm.mssm.edu/Enrichr/ |
| Seurat 3 | R. Satija Lab | https://satijalab.org/seurat/ |
